# Spinal cord associative plasticity improves forelimb sensorimotor function after cervical injury

**DOI:** 10.1101/2020.12.07.398289

**Authors:** Ajay Pal, HongGeun Park, Aditya Ramamurthy, Ahmet S. Asan, Thelma Bethea, Meenu Johnkutty, Jason B. Carmel

**Author notes:** Correspondence to: Jason B. Carmel Full address: Movement Recovery Lab, 650 West 168th Street Carroll Labs, Black Building 14th Floor Room 1412, New York, NY 10032, USA.

## Abstract

Associative plasticity occurs when two stimuli converge on a common neural target. Previous efforts to promote associative plasticity have targeted cortex, with variable and moderate effects. In addition, the targeted circuits are inferred, rather than tested directly. In contrast, we sought to target the strong convergence between motor and sensory systems in the spinal cord.

We developed spinal cord associative plasticity (SCAP), precisely timed pairing of motor cortex and dorsal spinal cord stimulations, to target this interaction. We tested the hypothesis that properly timed paired stimulation would strengthen the sensorimotor connections in the spinal cord and improve recovery after spinal cord injury (SCI). We tested physiological effects of paired stimulation, the pathways that mediate it, and its function in a preclinical trial.

Subthreshold spinal cord stimulation strongly augmented motor cortex evoked muscle potentials at the time they were paired, but only when they arrived synchronously in the spinal cord. This paired stimulation effect depended on both cortical descending motor and spinal cord proprioceptive afferents; selective inactivation of either of these pathways fully abrogated the paired stimulation effect. SCAP, repetitive pairing of these pathways for 5 or 30 minutes in awake rats, increased spinal excitability for hours after pairing ended. To apply SCAP as therapy, we optimized the parameters to promote strong and long-lasting effects. This effect was just as strong in rats with cervical SCI as in uninjured rats, demonstrating that spared connections after moderate SCI were sufficient to support plasticity. In a blinded trial, rats received a moderate C4 contusive SCI. Ten days after injury, they were randomized to 30 minutes of SCAP each day for 10 days or sham stimulation. Rats with SCAP had significantly improved function on the primary outcome measure, a test of dexterity during manipulation of food, at 50 days after SCI. In addition, rats with SCAP had persistently stronger responses to cortical and spinal stimulation than sham stimulation rats, indicating a spinal locus of plasticity. SCAP rats had lasting improvements in H-reflex modulation. The groups had no difference in the rat grimace scale, a measure of pain.

We conclude that SCAP strengthens sensorimotor connections within the spinal cord, resulting in improved reflex modulation and forelimb function after moderate SCI. Since both motor cortex and spinal cord stimulation are performed routinely in humans, this approach can be trialed in people with SCI or other disorders that damage sensorimotor connections and impair dexterity.

## Introduction

Integration of sensory feedback with motor commands is required for skilled movement. Neuromodulation strategies have been developed to target sensorimotor integration^1^. The most common application, paired associative stimulation, repetitively pairs median nerve stimulation with motor cortex stimulation timed to increase cortical excitability in healthy people^2^ and improve arm and hand function after injury^3, 4^. Using a similar logic but targeting the spinal cord, epidural stimulation has been used to drive activation of spinal afferents that can be timed to converge with descending motor activation. This strategy helped to restore walking after SCI, but only when spinal stimulation was timed to coincide with motor cortex signals arriving in the spinal cord^5–7^. Our goal was to employ associative learning through pairs of electrical stimuli to strengthen residual connections after cervical SCI, the most common type^8^. For people with cervical SCI, recovery of arm and hand function is the highest priority^9^.

Previous trials of paired stimulation targeting the spinal cord have targeted the motor system alone. Electrical stimulation of motor cortex has been paired with peripheral nerve stimulation to backfire spinal motoneurons. This motor system stimulation promotes lasting changes in motor evoked potentials and improved grip strength and finger control in people^10–12^. Intraspinal stimulation in monkeys has been timed with cortical activity so endogenous brain activity and exogenous spinal cord stimulation converge^13, 14^. Repetitive pairing induced a small but significant increase in evoked potentials.

In contrast to paired stimulation targeting cortex or the motor system in spinal cord, we targeted sensorimotor interactions in spinal cord with paired stimulation of motor cortex and cervical spinal cord^15^. The convergence of subthreshold spinal cord electrical stimulation with a descending cortical volley robustly augmented the cortical motor evoked potentials (MEPs). When this pairing was performed repeatedly, there was an increase in excitability that lasted for at least an hour. This plasticity required paired stimulation delivered at the proper time, but the exact neural circuits that mediate the sensorimotor interactions are not known. Knowing the targets is critical to ensure strong and selective engagement and to understand the anatomical basis for spinal cord associative plasticity (SCAP). It is also not known whether the sparse connections that are spared after SCI are sufficient to support SCAP and recovery of function. Together, the experiments in this study were designed to improve our understanding of SCAP mechanisms and to test its efficacy for the recovery of dexterity after SCI.

## Materials and methods

In awake and freely moving rats, we tested spinal cord associative plasticity (SCAP) and its efficacy on physiology and function in rats with SCI. We used custom spinal stimulating electrodes that can be inserted into the thin epidural space over the cervical spinal cord, are supple enough to conform to the spinal cord, and move with the neck^16, 17^. First, we tested the motor cortex and spinal cord stimulation timing with the hypothesis that convergence in the spinal cord would produce the largest changes in physiology. Second, we tested the necessity of corticospinal and segmental afferent connections for paired stimulation by selectively inactivating each using chemogenetic tools. We then paired motor cortex and spinal cord stimulation repeatedly to induce plasticity (SCAP). We optimized the protocol by varying the frequency, number, and pattern of stimuli in uninjured rats. We tested SCAP in rats with SCI. Finally, we performed a randomized, blinded preclinical trial of 10 days of SCAP versus sham stimulation to determine the effects on physiology and behavior after a moderate C4 contusion SCI. The primary outcome measure was a behavioral test of food-manipulation; several physiological, behavioral, and anatomical measures were secondary.

### Timing

In adult Sprague-Dawley female rats, we implanted three sets of electrodes: epidural screw electrodes over each motor cortex, EMG electrodes in each biceps muscle, and spinal epidural electrodes over the midline of dorsal C5-C6 spinal cord. To determine whether subthreshold spinal cord stimulation augments cortical motor evoked potentials (MEPs), we compared cortex stimulation only to motor cortex stimulation paired with spinal cord stimulation. We determined the threshold for provoking an MEP from motor cortex and from the spinal cord. We measured the time from motor cortex stimulation to the spinal cord dorsum potential (CDP) (Fig. 1A1). We then quantified the biceps MEP as the area under the curve in response to stimulation at 110% of threshold intensity (Fig. 1A2). In uninjured rats, this was compared against motor cortex stimulation paired with spinal cord stimulation at 90% of threshold intensity (Fig. 1A3 & 1B). This experiment quantified the size of the MEP with paired stimulation relative to cortex stimulation alone using unpaired t-tests with Bonferroni correction. We also tested whether stimulating motor cortex at 90% of threshold with spinal cord stimulation also at 90% would provoke an MEP together when neither produced a response on their own. The magnitude of the MEP was compared to that of motor cortex stimulation at 110% of threshold (Fig.1C). Finally, we tested the effects of paired stimulations in rats, weeks after a moderate C4 contusion injury (Fig.1D, using the same methods as Fig.1B but in rats with SCI).

**Figure 1.**
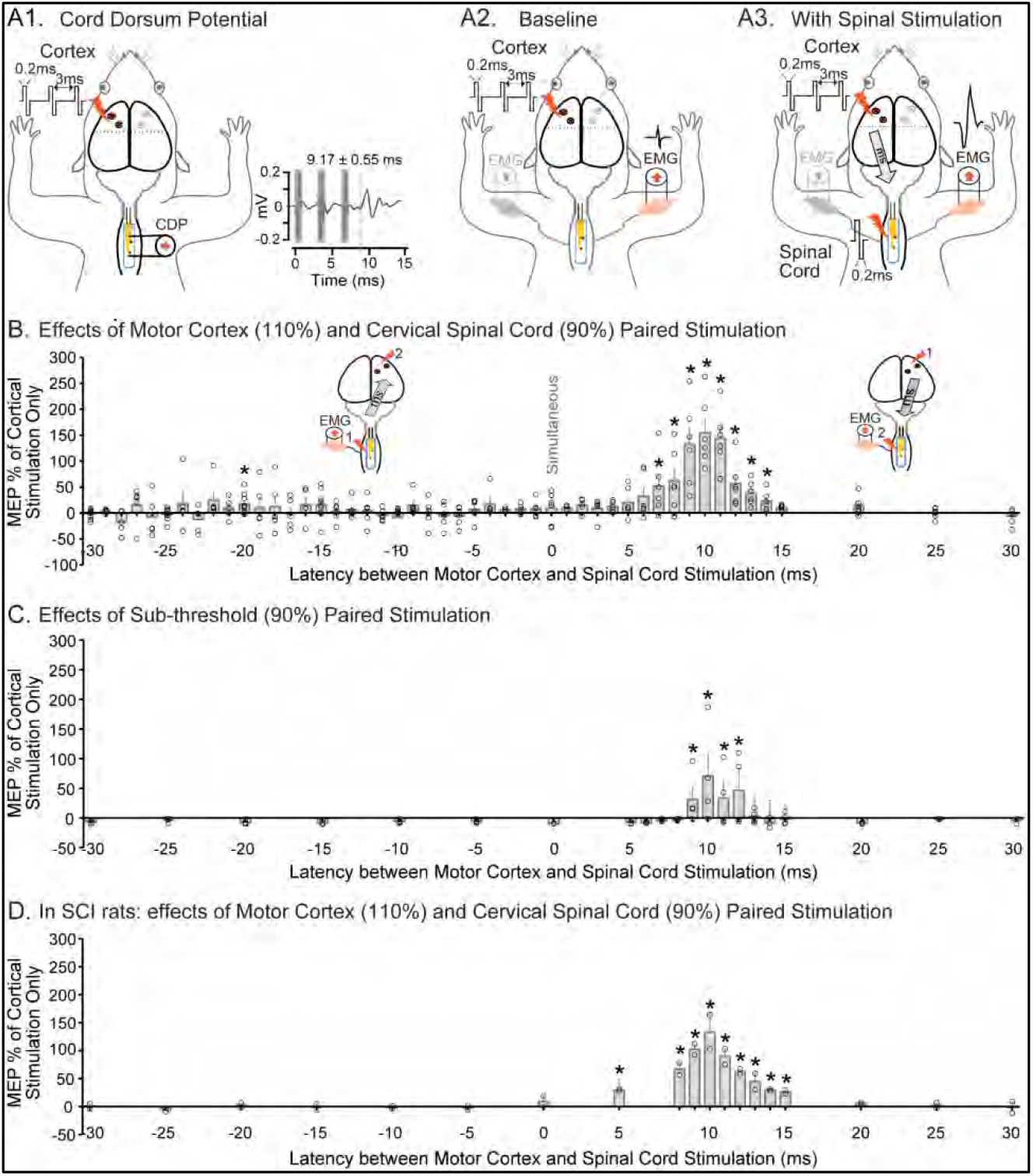
The immediate effects of pairing depend on timing. **(A1)** Schematic of cervical spinal cord dorsum potential (CDP) recording. After the onset of 3 pulses of motor cortex stimulation at 110% of motor threshold, the latency of the response was 9.17±0.55ms (*n*=3). (**A2).** Schematic of baseline testing after M1 stimulation at 110% of cortical threshold and EMG recorded from contralateral biceps. (**A3).** Immediate effect testing. Supra-threshold (110% of cortical threshold) stimulation of the motor cortex produces an MEP that is modulated by subthreshold (90% of spinal threshold) spinal cord stimulation. (**B).** Immediate effects of paired stimulation depend on latency. We tested different latencies between motor cortex and spinal cord stimulation from −30ms to 30ms. As indicated by the insets, negative times indicate that spinal cord was stimulated before motor cortex stimulation, and positive times indicate that spinal cord was stimulated after motor cortex stimulation. Maximum augmentation was found when spinal cord stimulation was delivered 10ms after motor cortex stimulation. MEPs were significantly elevated at the time points indicated, as measured by t-test with Bonferroni correction compared to no spinal stimulation baseline, asterisks represent p<0.05 (*n*=9). (**C).** Immediate effects when both motor cortex and spinal cord stimulations were paired with their 90% of threshold intensities. The EMG recordings of stimulation of motor cortex only or spinal cord only at 90% are shown in the insets. The paired stimulations with latencies between ±30ms were applied, and MEP responses were recorded. This figure shows % increase in the MEPs as a function of latencies. MEP responses started to appear with latencies around 10ms, and cortical convergence does not seem to cause any MEP response. MEP response becomes visible once the baseline EMG activity is doubled (*n*=4). (**D**). We further tested this hypothesis in animals with C4 spinal contusion. A moderate C4 contusion injury was performed in two rats, and spinal electrode arrays were inserted over C5-C6. Two weeks later, paired stimulation was performed. In SCI rats when 110% cortical stimulation were paired with 90% spinal stimulation, we observed a similar augmentation in MEP as observed in uninjured rats with peak at 10ms latency (134±20%, *p*=0.0001). Circle represents individual animals and error bars show SEM.

### Inactivation

To target the corticospinal tract (CST) selectively, a Cre-dependent adeno-associated virus (AAV1) encoding the Designer Receptor Exclusively Activated by Designer Drug (DREADD) was injected into the motor cortex and another encoding Cre-recombinase was injected into the cervical spinal cord (Fig. 2A1). Thus, only doubly infected neurons express the DREADD (Fig. 2A2), which can be inactivated by its ligand, clozapine-N-oxide (CNO)^18, 19^. One week after AAV injections, animals were trained on the vermicelli handling task^20, 21^. After reaching a stable baseline performance, the number of forepaw manipulations was measured before, during, and after inactivation (Fig. 2A3). One week after the behavioral test, rats were anesthetized and effects of inactivation on the immediate effects of paired motor cortex and spinal cord stimulation was tested (Fig. 2A4). Details in supplementary methods.

**Figure 2.**
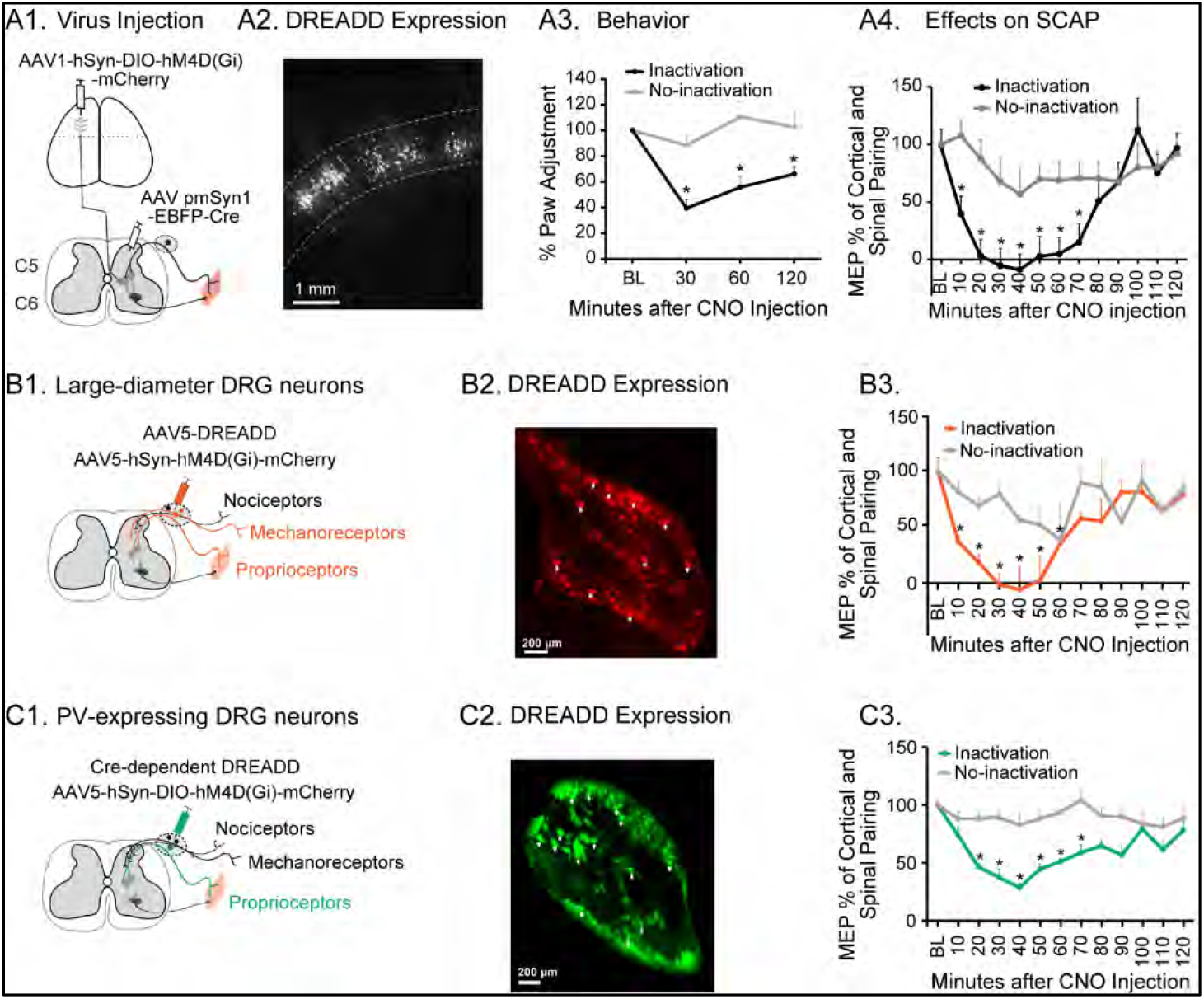
The corticospinal tract (CST) and spinal afferents are necessary for paired stimulation. (A) CST inactivation. **(A1)** AAV1-hSyn-DIO-hM4D(Gi)-mCherry was injected into the forelimb area of the left motor cortex and AAV2/retro-pmSyn1-EBFP-Cre was injected into the right side of the C5 & C6 spinal segments. (**A2)** Dual virus transduction drives robust mCherry expression in corticospinal neurons located in motor cortex layer 5 (dotted lines). (**A3)** CST inactivation impairs skilled forepaw use. The number of adjustments of both forepaws was counted while the animal was eating uncooked pasta before and at 30, 60, and 120 minutes after clozapine N-oxide (CNO) injection. Inactivation significantly reduces the paw manipulations in the paw controlled by the inactivated CST (*n*=3). (**A4)** CST inactivation abrogates the effect of paired stimulation. One week after the behavioral test, the CST was inactivated by CNO injection. The augmentation of paired stimulation was fully abrogated at 20-60 minutes in the inactivation side and gradually recovered (*n*=3). (**B**) **Large-diameter afferent inactivation**. (**B1)** An AAV5 encoding the inactivating DREADD was injected into DRG on same site as the targeted side. (**B2)** DREADD expression in neurons (arrowhead), infected with the virus measured 67.9±18.5µm in diameter, similar to neurons that mediate touch, proprioception, and muscle length/tension. (**B3)** Inactivation abrogates paired stimulation in rats infected with AAV5-DREADD, inactivation side (red line) compared to non-inactivated side (grey line, *n*=2), 100% suppression at 40 minutes after CNO injection. **(C) Proprioceptor inactivation.** (**C1)** A Cre-dependent DREADD was injected into the DRG on the inactivation side of rats expressing Cre under control of parvalbumin, which is expressed selectively by proprioceptors. (**C2)** DREADD in proprioceptive neurons (arrowhead), infected with the virus. **C3.** Inactivation abrogates paired stimulation in PV-Cre rats, inactivation side (green line) compared to baseline (*n*=3); more than 70% suppression of paired stimulation effect was observed at 40 minutes after CNO injection. More selective inactivation (green line) resulted in 25.8%±3.8% suppression of sensory afferents while non-selective (red line; −1.5%±10.5%) inactivation caused more robust suppression of paired stimulation responses on the targeted side of forelimb after CNO injection. Vehicle (normal saline) injection did not change MEPs compared to baseline on the inactivated side (data shown in supplemental Fig.1). Error bars show SEM.

The necessity of afferents for the immediate effects of paired stimulation was tested using two inactivation strategies. For the first strategy, large diameter dorsal root ganglion (DRG) neurons were inactivated (Fig. 2B1). Large DRG neurons mediate touch, proprioception, and muscle length/tension^22^. The specificity of large-diameter DRG neurons was provided by the tropism of the AAV5^23^. An AAV5-hSyn-hM4D(Gi) was injected into each DRG at C5, C6, and C7 on one side of the spinal cord after partial laminectomy (Fig. 2B2). At the time of virus injection, cortical, EMG and spinal electrodes were implanted as described earlier. CNO was administered as described in the supplementary methods, and responses to paired stimulation was measured (Fig. 2B3).

For the second strategy, we targeted the proprioceptive afferents, since these mediate the effects of spinal cord stimulation (Fig. 2C1)^24–26^. We used a genetic strategy to target DRG neurons that express parvalbumin, which are proprioceptors^27, 28^. In transgenic rats that express Cre recombinase under the control of parvalbumin promoter PV-Cre rats (LE-Tg(Pvalb-iCre)2Ottc), we injected a Cre-dependent DREADD (AAV5-hSyn-DIO-hM4D(Gi)-mCherry^29^ into the C5, C6, and C7 DRGs as described above (Fig. 2C2). Inactivation with CNO and physiology measurement were also identical (Fig. 2C3).

### Repetitive pairing (SCAP)

We tested brain and spinal cord responses at baseline and then after the period of repetitive pairing (Fig. 3A1-A3), as we did in Mishra et al.,^15^ but in awake rats with implanted electrodes. This differs from the immediate effects of stimulation (Fig. 1A3), as we examined the lasting effects of paired stimulation. Before modulation, we measured the stimulation threshold for both motor cortex and spinal cord, defined as the stimulation intensity needed to produce an MEP in 50% of trials. We then stimulated with 110% of threshold to determine cortical and spinal MEPs, which were quantified as the area under the curve. At baseline, cortical and spinal MEPs were measured singly and independently (Fig. 3A1). For repetitive pairing (Fig. 3A2), motor cortex stimulation was performed at 110% of the threshold for evoking a cortical MEP followed 10ms later by spinal cord stimulation performed at 90% of the threshold for generating a spinal MEP. After repetitive pairing (Fig. 3A3), the same measures at baseline were taken immediately after pairing and every 20min thereafter up to 120min. We quantified excitability as 1/threshold and expressed it as % change from baseline.

**Figure 3.**
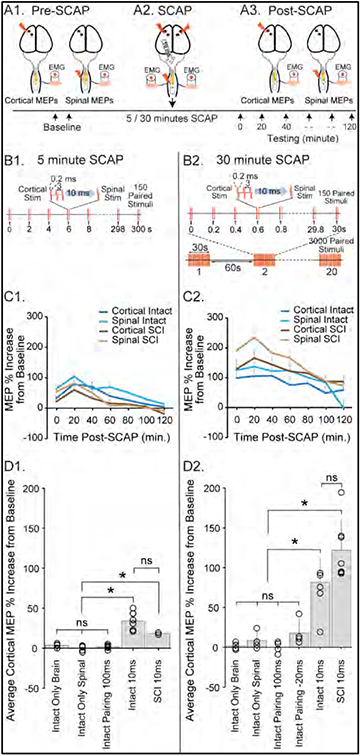
Spinal cord associative plasticity (SCAP) protocol. **(A1)** Baseline measurements of cortical MEP, spinal MEPs and thresholds before repetitive pairing. Cortical and spinal stimulation at 110% of the cortical and spinal thresholds. MEPs recorded from the biceps. Threshold was determined by adjusting stimulation intensity to provoking a short-latency EMG in >50% of trials. (**A2)** Repetitive paired stimulation, repetitive pairing motor cortex and spinal cord stimulation at 10ms latency for either 5 or 30-minutes. (**A3)** Testing the effects of both 5 and 30 minutes SCAP, cortical, spinal MEPs and thresholds were measured as it was done before SCAP. (**B1)** 5-minute SCAP protocol (0.5 Hz, 150 pairs). (**B2)** A 30-minutes SCAP paradigm (5 Hz, 150 pairs of cortical and spinal stimuli 20 times with 1-minute gap between bursts, delivering a total of 3000 pairs). (**C1 – D2)** SCAP causes lasting changes in cortical and spinal cord MEPs in uninjured rats and rats with SCI. (**C1)** Cortical MEP augmentation after 5-minute SCAP in uninjured awake rats (*n*=6) as well as in rats with C4 SCI (*n*=2). In both injured and uninjured groups of rats after 5-minutes of repetitive pairing, cortical evoked responses were augmented for more than 1 hours and this augmentation persisted even after SCI, then returned to the baseline. A similar effect was also observed with the spinal MEP and excitability increased after pairing (data shown in S. Fig. 4A). (**C2)** Cortical MEP augmentation after 30-minute SCAP in uninjured awake rats (*n*=4) and in SCI rats (*n*=7). In both SCI and intact (uninjured) groups of rats after 30-minutes of repetitive pairing, cortical evoked responses were augmented for more than 2 hours and this augmentation persisted even after SCI, similarly the spinal MEP and excitability increased for 2 hours after pairing (data shown in S. Fig. 4B). **(D1)** Average MEP augmentation over 120 minutes with 5-minutes SCAP conditions. Only motor cortex (4%±7%) or only spinal cord stimulation (8%±6%) or pairing at an inappropriate latency (100ms; 2%±5%), do not result in elevated MEP responses. (**D2)** Average MEP augmentation over 120 minutes with different 30-min SCAP conditions. Only motor cortex (1.3%±6%) or only spinal cord stimulation (8%±4%) or pairing at an inappropriate latency (100ms; 0.2%±3%), do not result in elevated MEP responses. Pairing at −20ms aiming at cortical convergence, does not cause a strong MEP response (18%±5%). Circle represents individual animals and error bars shows SEM. A One-way ANOVA was done for overall comparison and was highly significant followed by Bonferroni correction for multiple comparisons. * indicates *p*<0.001, ns=non-significant.

### Optimization

In order to maximize the efficacy of the SCAP protocol (5-minute SCAP, Fig 3B1), we varied four stimulation parameters: frequency, number of stimuli, number of trains, and time between trains. We kept the total stimulation period to 30 minutes, a period that could be used for stimulation in people. In two rats implanted with cortical and spinal electrodes, we first varied frequency by delivering 10 stimuli at 5Hz, 2Hz, 1Hz, 0.5Hz and 0.2Hz and measured the size of the MEPs (Supplementary Fig. 3A). For the other parameters, we measured the change in cortical MEPs and cortical and spinal excitability. Spinal excitability was used since this was less variable than spinal MEPs, and out of concern that eliciting spinal MEPs might induce plasticity. For these trials, we used 5 Hz pairing and varied the number of stimuli by comparing 150, 75 & 38 pulses (Supplementary Fig. 3B, C & D1-D3). For subsequent trials, we used 150 stimuli, which we term a train, and varied the number of trains, either 10 or 20 (Supplementary Fig. 3B, C & F1-F3). For subsequent trials, we used 20 trains and varied the time between trains, either 1 or 5 minutes (Supplementary Fig. 3B, C & E1-E3).

### Cervical spinal cord contusion (SCI)

The C4 spinal cord contusion surgery was performed as previously described^30^. Briefly, the C3 to C5 vertebrae were exposed by skin incision and muscle separation, and laminectomy of C4 was performed. An Infinite Horizon impactor (IH-0400, Precision Systems & Instrumentation, LLC), was used to deliver 200 kilodynes force onto the spinal cord with a 3.5mm diameter tip. The force curve of the impactor was recorded (Fig. 7A). After the injury, the spinal electrode array was implanted as described above and rats received intense care, as described in the supplementary methods.

### Preclinical trial

We designed a preclinical efficacy trial; the timeline is shown in Fig. 4A. Ten days after SCI, rats were randomized to receive real or sham SCAP for 10 days, a period that we have previously shown to be therapeutic^30, 31^. The daily stimulation duration (30 minutes) and the number of sessions (10) were designed to be clinically viable. The primary outcome measure was the dexterity (Irvine, Beatties, and Bresnahan scale^32^) score, and secondary outcomes included performance of skilled walking, changes in MEPs (immediate effects), thresholds, lasting effects of pairing in stimulated rats, H-reflex modulation (a measure of hyperreflexia), and a pain measure (RGS). The study was powered based on the effect size we observe of a different neuromodulation strategy^30^. A priori power analysis suggested we need 8 rats in each group to achieve a power of 0.8. Rats were pseudorandomized into real (Stim) and sham (Control) groups with each cohort of up to 8 rats having rats assigned to each group. Rats were randomized immediately after SCI surgery, and experiments were carried out with blinded measures.

**Figure 4.**
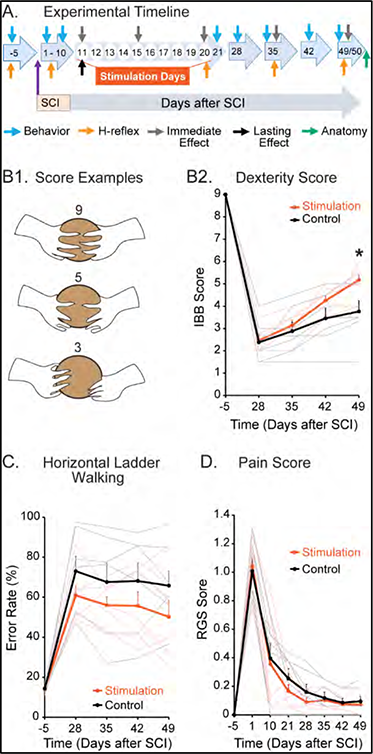
SCAP promotes motor recovery after SCI and does not affect pain. **(A)** Experimental timeline: Rats were trained on the food manipulation task and then subjected to C4 contusion injury. Rats were randomized to SCAP daily for 10 days (stimulation group) versus sham (no) stimulation (control group). Physiology and behavior testing was done before stimulation on and then weekly after stimulation. After 50 days, animals were sacrificed, and anatomical analysis was done. (**B1)** Schematics showing characteristic features of rat forelimbs during cereal manipulation task, the primary outcome measure. Images of a representative stimulation rat were generated from frames of live videos with the image tracing tool of Adobe Illustrator software. Score 9 at baseline before SCI: Volar (palm) support, paw adapts to the shape of the cereal. Score 5 after stimulation: Volar support, digit 2 contributes to cereal manipulation, other digits extended. Score 3 after injury, before stimulation: Lack of volar support, extended digits, and paw doesn’t adapt to the shape of the cereal. (**B2)** Average and individual rat’s IBB scores. At each time point, the IBB score was averaged over four pieces of cereals for cereal manipulation trials. Thick red and black line are the average IBB scores for all stimulation group rats and controls respectively. Thin light red and black lines indicate individual stimulation and control group rats respectively. Error bar represents SEM. * unpaired t-test, *p*=0.024, at 49 days after SCI (*n*=7 in both stimulation and control groups). (**C)** Locomotor behavior on horizontal ladder shows % error that rats made while walking across the horizontal ladder over 20–24 trials at each time point, averaged over left and right forelimbs. Thick red and black lines are the average error rate for all stimulation group rats and controls respectively. Thin light red and black lines indicate individual stimulation and control group rats respectively. There was a decreased trend in % error rate in the rats in stimulation groups as compared to controls but was not significantly deferent between the groups at the final end point of testing (day 49, unpaired t-test, *p*=0.182, *n*=7 in each group). **(D)** Pain scores quantified pain using the rats facial characteristics. Thick red and black lines are the average pain scores for all stimulation group rats and controls respectively. Thin light red and black lines indicate individual stimulation and control group rats respectively. We observed no increase in pain score with 10 days of repetitive stimulation, rats in both the groups did not experience pain during the stimulation phase (unpaired t-test, *p*=0.646, *n*=7 in each group). Error bars show SEM.

### Behavior: Food Manipulation Task

To assess the rat’s ability to manipulate, we used the Irvine, Beatties, and Bresnahan scale (IBB;^32, 33^, as we did previously^30^. **Horizontal ladder walking task**: The horizontal ladder walking task measured paw placement accuracy on irregularly spaced rungs, a task that requires the corticospinal tract^31, 34–36^. **Rat Grimace Scale (RGS):** This scale is used to quantify pain based on rat facial characteristics in a way that is commonly used to assess pain in people^37^.

### Physiology

The immediate effects of pairing (Fig. 1A3) as well as thresholds were tested before the 30-minute SCAP on each day of stimulation and every two weeks thereafter. The lasting effect of SCAP (Fig. 3A1) was measured on day 11 after SCI.

### H-reflex testing

To track H-reflex modulation, we measured the H-reflex at baseline, one week after SCI, just after 10 days of stimulation, and every two weeks post-stimulation (Fig. 4A). First, we tested responses to increasing stimulus intensities up to 2X threshold, which activates low threshold afferent fibers. Next, we tested the frequency-dependent depression of the H-reflex^38^.

### Anatomy

To determine the severity of SCI, we measured impact force during spinal cord contusion. We quantified the volume of spared gray and white matter at the injury site. We also quantified spared axons below the lesion in both groups. For CST axon quantification, BDA was injected into cortex and histology was performed on sections above (C3) and below (C7) the lesion site as in our previous studies^30^. For sensory axon quantifications, AAV1 was injected in C5-C7 DRG, and sensory afferent axons were quantified below the lesion segments.

### Statistical Analyses

All the comparisons between stimulation conditions were done with paired/ t-tests within group and unpaired t-tests between groups with Bonferroni correction for single time points or one-way ANOVA with Bonferroni corrections for multiple time points. Cohen’s d was used effect size. All the statistical analysis was performed with SPSS (SPSS Statistics for Windows, version 27; IBM Corp., Armonk, NY, USA). We tested the normality of the data using the Shapiro-Wilk test. All data are expressed as Mean ± SEM.

### Data Availability

The datasets generated and analyzed during the current study are available from the corresponding author on reasonable request.

## Results

We hypothesized that repetitive pairing of motor cortex and spinal cord would augment spinal excitability and improve function in rats with SCI through spinal sensory and descending motor convergence in the spinal cord. This hypothesis was tested in steps. First, we tested whether convergence of paired stimulation was more effective in cortex or the spinal cord by altering the timing for paired stimulation. Second, we tested whether the connections preserved by cervical SCI were sufficient to enable paired stimulation. Third, to determine whether the CST and proprioceptors were necessary, we selectively inactivated each. Fourth, we optimized the repetitive application of paired stimulation (SCAP) protocol. Fifth, we tested this optimized protocol in rats with SCI. Finally, we tested whether 10 days of repetitive SCAP improved physiology and behavior after SCI.

### Spinal stimulation augments cortical responses when properly timed

We hypothesized that stimulating the spinal cord at the time that descending motor potentials arrive would most strongly augment cortical MEPs. The latencies shown in Fig. 1A1 are the averages of 20 responses each from three animals, while the waveforms are for a representative trial in one animal. As shown in Fig. 1A1, recorded C5–C6 spinal cord responses to cortical stimulation in awake rats. After the initiation of a train of 3 stimuli in the motor cortex, the CDP arrived at 9.17±0.55ms (Mean±SEM, n=3). Spinal cord stimulation augmented cortical MEPs at multiple time points (Fig. 1B) with a strong peak when the spinal cord was stimulated 10ms after the motor cortex (155%±27%, *df*=13, *Cohen’s d*=40.86, *p*<0.0001). There were also smaller increases in MEPs when the spinal cord was stimulated 20ms before cortex (17%±6%, *df*=16, *Cohen’s d*=13.50, p=0.019).

If the effects of motor cortex and spinal cord stimulation are synergistic, then subthreshold stimulation at each site may result in a suprathreshold response if they are properly timed. To test this hypothesis, stimulation intensities for both cortical and spinal stimulations were set to 90% of the threshold. Stimulation of the spinal cord only and motor cortex only produced no responses. Paired subthreshold stimulation generated an MEP with a peak effect at 10ms. The response was larger than cortical stimulation only at 110% of threshold (66±38% larger) of 110% cortical threshold, p=0.001; paired t-test). Thus, the synergistic effects of subthreshold stimulation were observed only with convergence in the spinal cord.

### Neural circuits spared by SCI are sufficient to mediate the effects of paired stimulation

To determine if the neural circuits spared by SCI were sufficient to enable paired stimulation, we tested paired stimulation in rats with moderate C4 spinal contusion. Just as in Fig. 1A, 110% cortical stimulation was paired with 90% spinal stimulation. Augmentation of MEPs was observed with a peak at 10ms latency (Fig. 1D; 134±20%, p=0.0001, paired t-test). No augmentation was observed at any latency in which spinal cord was stimulated before cortex. This supports the hypothesis that the spinal cord pathways spared by contusion can mediate the effects of sensory-motor convergence in the spinal cord.

### Effects of paired stimulations are mediated by corticospinal and proprioceptive afferents

To test the hypothesis that descending corticospinal circuits and large-diameter afferents are necessary for effects of pairing, we selectively inactivated each with a chemogenetic tool. In rats with injections into motor cortex and spinal cord (Fig. 2A1), layer 5 neurons in the motor cortex were selectively and robustly labeled (Fig. 2A2). These neurons were inactivated by administration of CNO, the ligand for the inhibitory DREADD. At the time of maximum inactivation (30min; Fig. 2A3) the number of manipulations the rat made while eating uncooked pasta with the paw targeted for inactivation diminished to 39%*±7*%. The number of manipulations partially recovered by 120 minutes, as the CNO washed out. Manipulations of the non-targeted paw were not highly affected. We then tested whether this inactivation abrogated the effects of paired stimulation (Fig. 2A4). At baseline, subthreshold spinal cord stimulation delivered 10ms after suprathreshold motor cortex stimulation strongly augmented biceps MEPs (>100%). Administration of CNO caused the MEP augmentation to be diminished compared to baseline, and at 30 minutes (−6.3% ±23%, *df* =4, *Cohen’s d*=5.35, *p*=0.003), spinal cord stimulation had no augmenting effect on motor cortex MEPs on the inactivated side. The effects of pairing recovered by the end of the 120-minute testing period. On the non-inactivated side, there was a less pronounced and more delayed effect of inactivation. Thus, corticospinal neurons are necessary for the effects of paired stimulation.

To determine if paired stimulation also depends on spinal afferents, we inactivated dorsal root ganglia (DRG) neurons using two methods. First, we targeted large-diameter neurons; an AAV5 encoding an inactivating DREADD was injected into DRGs of C5 and C6 on one side (Figure 2B1). Cells infected with the virus (Fig. 2B2) measured 68±18µm in diameter, like neurons that mediate touch, proprioception, and muscle length or tension^22^. Before inactivation, the pairing of the motor cortex and spinal cord stimulation caused a large augmentation of the biceps MEP (>100%; Fig. 2B3, red) compared to cortical stimulation only. Thirty minutes after CNO injection, the paired stimulation effect was abolished (−1.5%±10.5%, *df*=4, *Cohen’s d*=2.34, *p*=0.013) and largely returned by 120 minutes. There was no significant effect on the side without inactivation.

For the second approach, we selectively targeted the proprioceptors. A Cre-dependent DREADD was injected into the DRGs (Fig. 2C1) of transgenic rats expressing Cre under control of parvalbumin, which is expressed selectively by proprioceptors (Fig. 2C2). Inactivation caused a large-scale, but incomplete, abrogation of paired stimulation (*28.6%±3.8%, df=4, Cohen’s d=11.81, p=0.017*; Fig. 2C3, green). Inactivation also decreases the motor cortex excitability (supplementary fig. 1A), spinal MEPs (supplementary fig. 1B), and spinal excitability (supplementary fig. 1C) in PV-Cre rats. The loss of paired stimulation effect with inactivation of proprioceptors was less than inactivation of many large-diameter DRG neurons (including mechanoreceptors and others). Thus, large-diameter sensory afferents, and proprioceptors in particular, are necessary for the effects of SCAP.

To ensure that the effects of inactivation were due to loss of function of specific circuits, we conducted two control experiments. We tested whether CNO injection itself changed physiological responses. In a blinded test of rats without viral injection, CNO had no effect on physiological responses (supplementary fig. 2A). In addition, administration of saline in rats with DREADD injection showed no change in paired stimulation effects on cortical MEPs, cortical excitability, spinal MEPs, or spinal excitability (supplementary fig. 2B1-4). Thus, neither CNO without DREADD administration nor DREADD injection without CNO affected physiological responses.

### SCAP caused lasting changes in cortical and spinal cord evoked responses in awake rats with and without SCI

In addition to the immediate effects of paired stimulation, repeatedly pairing motor cortex and cervical spinal cord stimulation at 10ms latency promotes spinal cord associative plasticity (SCAP). In these experiments, cortical and spinal responses were tested before and after a period of SCAP (Fig. 3A). SCAP delivered over 5-minutes (3B1) nearly doubled the size of both the cortical and spinal MEPs (3C1, dark and light brown lines), an effect that ebbed over 120 minutes. These changes were of a similar magnitude but much longer duration than we previously reported; anesthetized rats had no increase in MEPs after 60 minutes^15^.

We also tested the effects of SCAP in rats with SCI. Following the 5-minute SCAP protocol, rats with SCI also showed lasting increases in the size of cortical and spinal MEPs (Fig. 3C1, dark and light blue lines). Similar effects were also observed with cortical and spinal thresholds in intact and SCI rats (Supplementary Fig. 4A). We compared the average effects of paired stimulation delivered at 10ms latency versus several control conditions: motor cortex stimulation alone, spinal cord stimulation alone, pairing at 100ms latency, and 10ms pairing in intact and SCI rats (3D1). The control conditions did not induce plasticity (AVOVA with Bonferroni correction; p=0.65). However, both intact and SCI rats had significant increased cortical MEPs compared with these control conditions (AVOVA with Bonferroni correction p<0.001); there was no significant difference between intact rats and rats with SCI (Intact=38%±11%; SCI=18%±10%; AVOVA with Bonferroni correction; p=0.11). The change in cortical MEP was averaged across the 120 minutes of testing. Thus, the circuits spared by cervical SCI were sufficient to enable SCAP.

In anticipation of using SCAP in a preclinical therapeutic trial, we optimized the protocol. We kept the duration to a maximum of 30 minutes, a period that is tolerable for clinical neuromodulation^39^. We tested 4 parameters: frequency of pairing, pulse number (length of train), time between trains and number of bursts/trains (supplementary Fig. 3). We tested five frequencies to determine if each pair of stimuli would cause inhibition or augmentation of subsequent stimuli. The 5Hz and 0.2Hz frequencies produced little change between stimuli, and we selected 5Hz to minimize the total duration of stimulation. We also observed that 1Hz was strongly inhibitory. Using the 5Hz frequency, we varied the number of stimuli; 150 stimuli were better than the others tested. We tested the time interval between bursts of 150 stimuli at 5 Hz; we selected 1 minute to balance effect size and the total duration of stimulation. Finally, we tested the total number of bursts. We observed that 20 trains of stimulation had a more durable effect than 10 trains with a similar overall magnitude. This resulted in the paradigm pictured in Figure 3B2 which delivers 3000 pairs of stimuli over 30 minutes.

We tested the effects of 30-minute SCAP in intact rats and rats with SCI. In intact rats, cortical and spinal MEPs were doubled immediately after 30-minute SCAP (Fig. 3C2, dark and light blue lines), and they were still ∼50% elevated at 120 minutes. In SCI rats, following the 30-minute SCAP protocol (3B2) the size of both the cortical and spinal MEPs was increased 100-200% (3C2, dark and light brown lines), and the MEPs were still almost double at 120 minutes. Similar effects were also observed with cortical and spinal excitability in both intact rats and those with SCI (supplementary fig. 4B). The physiological effects of the 30-minute SCAP paradigm are more robust than the 5-minute SCAP paradigm in both intact and injured animals. Further, we compared the efficacy of 30-minute SCAP to control conditions (motor cortex alone, spinal cord alone, or paired with spinal cord stimulation after a 100 ms delay) and convergence in motor cortex. Neither the control conditions nor convergence at cortex increased cortical or spinal MEPs (AVOVA with Bonferroni correction; p=0.20; Fig. 3D2) or changed cortical or spinal thresholds (supplementary Fig. 5). However, both intact and SCI rats had a significant increased cortical MEPs after 30-minute SCAP compared with these control conditions (AVOVA with Bonferroni correction p<0.0001; Fig 3D2). There was no significant difference between intact rats and rats with SCI (Intact=82%±9%; SCI=121%±11%; AVOVA with Bonferroni correction; p=0.15). Finally, the similar magnitude and duration of effects in intact and injured rats suggest that circuits spared by SCI are sufficient to mediate pairing effects.

**Figure 5.**
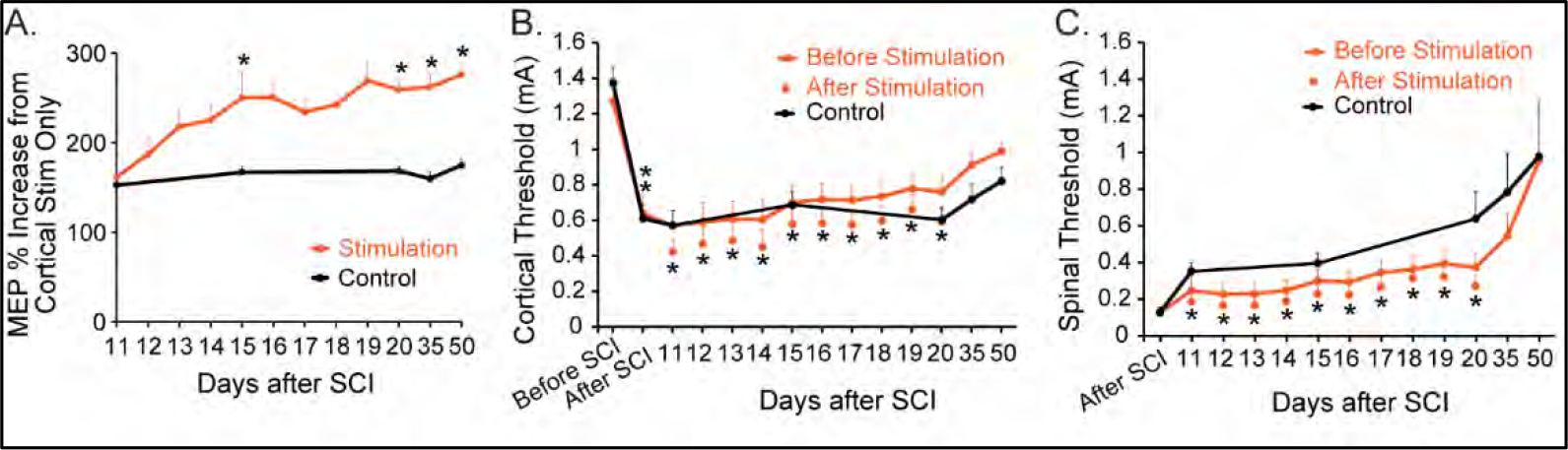
Ten days of SCAP produces augmented MEPs for at least 50 days after SCI. (**A)** The immediate effects were not different before stimulation (day 11) in between the stimulation and control groups, in the stimulation group (*n*=7), cortical MEP augmentation increased over the 10-day periods of stimulation to 259±12%, while the control group (*n*=7) did not change (168% ±5.4%). The difference between groups was maintained at 50 days after SCI (Stim=258%±21% vs Control=169%±10%, *df*=12, *Cohen’s d*=3.10, *p*=0.0001). This indicates that repetitive pairing over a 10-day period creates a durable change in physiology. (**B)** Cortical thresholds before SCI were above 1mA in both stimulation (red line) as well as control (black line) rats, after SCI there was a 50% drop in the cortical thresholds in both groups of rats, this may be due to hyperreflexia. There was gradual increase in cortical threshold after SCI but still below the before SCI value. The red line indicated the change in cortical thresholds in each session of repetitive paired stimulation (30-minute SCAP). After each session of 30-minutes of SCAP the cortical threshold decreased (red dots) as compared to that of the pre-stimulation value, indicating increased cortical excitability after each session. (**C)** Spinal thresholds after SCI were low in both stimulation (red line) as well as control (black line) rats, the spinal threshold was measured only after SCI since spinal electrodes were implanted just after spinal contusion. There was a gradual increase in spinal threshold after SCI that we observed with our spinal electrodes due to an increase in impedance post implantation. The red line indicates the change in the spinal thresholds in each session of repetitive paired stimulation (30-minute SCAP). After each session of 30 minutes of SCAP the spinal threshold decreases (red dots) as compared to that of pre-stimulation value, indicating increase spinal excitability after each session. Error bars show SEM.

### Repetitive SCAP after SCI improved forelimb dexterity without changing affective pain

To determine whether SCAP might be used to restore sensorimotor function after SCI, we performed a preclinical efficacy trial (Fig. 4A). The pre specified primary outcome measure was the IBB test, a scale that measures paw function while eating pieces of cereal (Fig. 4B1). Before SCI, all rats performed at the top of the 9-point scale (Fig. 4B2). Twenty-eight days after SCI, the rats in each group showed severe impairment (stimulation=2.46±21; control=2.38±34), with difficulty holding and manipulating the cereal. By 50 days after injury, the last time tested, control rats scored 3.76±0.49, and rats with stimulation scored 5.18±0.25 (50days; *df*=12, *Cohen’s d*=2.62, *p*=0.024) demonstrating better grasp and support of the cereal (Fig. 4B2).

Our secondary behavioral outcomes were walking performance on the horizontal ladder (4C) and observed pain (4D). Rats performed equally before the injury. For walking, by 28 days, the first time that rats reliably perform the task, control rats made 73%±7% errors, while rats with stimulation made 61%±5% errors. At the end of the testing period, control rats made 66%±8% errors, and rats with stimulation made 50%±8% errors; this difference was not significant (50 days: *df*=12, *p*=0.18). To test the effects of stimulation on pain, we used the RGS, which uses facial features to rate severity on a three-point scale. All rats exhibited pain after SCI (Fig. 4D), despite being given analgesics, which diminished by 10 days and stayed low. There was no difference in pain scores between the groups at the end of the trial (50 days: *df*=9, *p*=0.65). Thus, ten days of SCAP improved dexterity and did not produce pain after SCI.

### SCAP caused lasting increases in evoked responses and improved H-reflex modulation over weeks

We hypothesized that SCAP delivered daily over 10 days in rats with SCI would strengthen sensorimotor connections, as measured by evoked responses. We measured the immediate effect of paired stimulation-; how strongly spinal stimulation augmented the cortical MEP as the time they are paired. Since paired MEPs are always compared to cortical stimulation alone, this measure is stable and reflects the strength of convergent stimuli. In rats with SCI, the immediate effect of pairing was large at 11 days after SCI (Stim=162%±22%, Control=153%±11%) in both groups before the start of stimulation (Fig. 5A). In the stimulation group, cortical MEPs increased over the 10-day period to 259%±12%, while the control group did not change (168%±5.4%). The difference between groups was maintained at 50 days after SCI (Stim= 258%±21% vs Control = 169%±10%, *df*=12, *Cohen’s d*=3.10, *p*=0.0001). This indicates that repetitive pairing over a 10-day period creates a durable change in MEPs from paired stimulation.

We also measured cortical and spinal thresholds over the study. Cortical thresholds before SCI were above 1.2 mA in both stimulation (red line) and control (black line) rats (Fig. 5B). After SCI there was ∼50% drop in the cortical thresholds in both groups of rats (ANOVA, p<0.001). After each 30-minute SCAP session, the cortical threshold (red dot) was lower than when it began (AVOVA, p<0.001; red line). There was a gradual increase in cortical threshold until the end of the study period. Similarly, the spinal thresholds (Fig. 5C) immediately after SCI were low in both stimulation (red line) as well as control (black line) rats (electrodes were implanted just after SCI). There was a lower threshold in the rats with stimulation compared to control rats (AVOVA, p<0.001, red line vs black line). There was a gradual increase in spinal threshold over the study period, as we have observed in implanted arrays^16^.

We hypothesized that repetitive stimulation would help restore spinal reflex modulation. We tested H-reflex modulation before and after SCI (Fig. 6A). H-reflexes were elicited at different interstimulus intervals to modulate responses (Fig. 6B). Before injury in both rats with SCAP (Fig. 6C) and controls (Fig. 6D), shorter intervals between H-reflexes caused the second response to be diminished (blue lines). After SCI, the rate-dependent decrease in the H-reflex was diminished (top lines). Ten days of SCAP largely restored rate-dependent depression of the H-reflex (6C; bright red lines). In contrast, control rats had little to no recovery of H-reflex modulation (6D; gray lines). This effect was still persistent at 50 days after SCI (control 63%±1.5% vs Stim 45%±1.7%; *df*=12, *Cohen’s d*=4.33, *p*=0.0001). Thus, SCAP improved H-reflex modulation after SCI.

**Figure 6.**
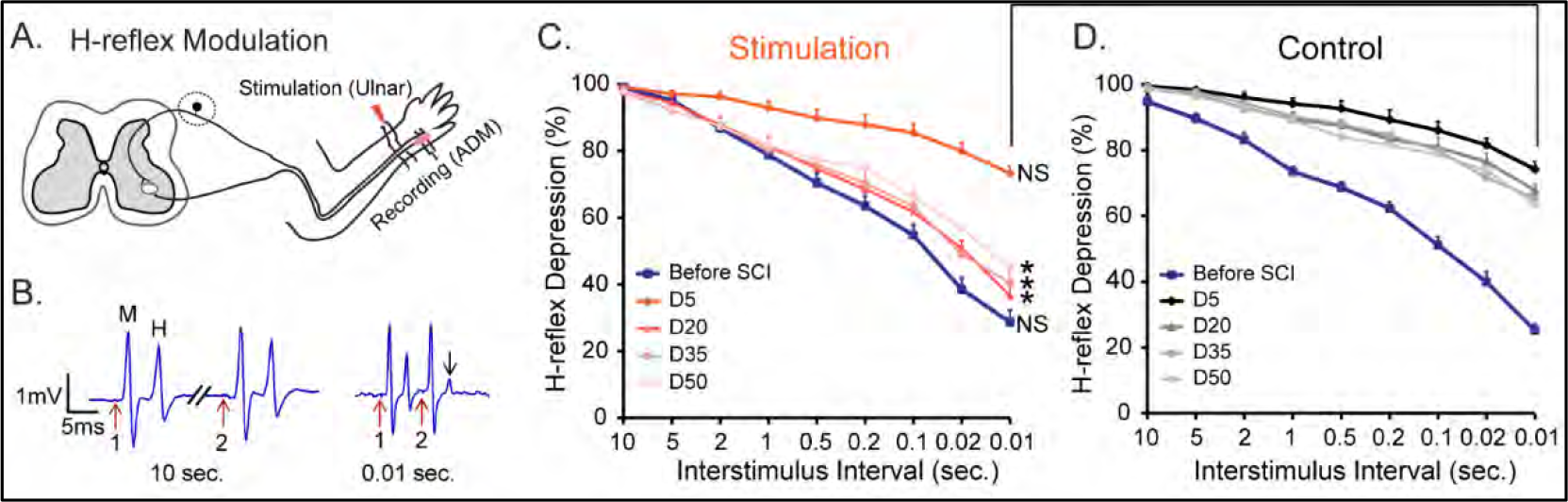
Improved H-reflex modulation with SCAP. **(A)** Method of H-reflex testing. (**B)** The second H-reflex is diminished (arrow) at short inter-stimulus intervals. (**C)** Rats with stimulation (*n*=7). Less H-reflex depression in rats 5 days after SCI (dark red) compared to before (blue). After paired stimulation H-reflex depression was largely restored and maintained in the stimulation group. (**D)** Control rats (*n*=7) do not show restoration of H-reflex modulation. M=M-wave, H=H-reflex. Error bars show SEM.

### No difference in SCI and tissue sparing between stimulation and control groups

To determine severity of SCI, we quantified the peak force rats received during C4 spinal contusion and the tissue preserved at the injury site at the end of the trial. The peak force that rats received is shown in Fig. 7A. There was no significant difference between the stimulation group (203.57±5.81 kilodynes) and the control group (209.1±7.13 kilodynes; t-test, *p*=0.28). We also compared the amount of tissue spared at the site of injury. Representative spinal cord sections are shown in Fig. 7B and representative lesion reconstruction (Fig. 7C, red). There was no significant difference in tissue sparing at the lesion epicenter between the control (3.13±0.46 mm^2^) and the stimulation group (2.76±0.53 mm^2^, independent t-test, *p*=0.61; Fig. 7D). We concluded that the injuries rats received are similar between stimulation and control groups. We quantified the length of the spared descending motor and sensory afferent axons below the lesion in both groups. We did not observe a significant difference in total axon length of both descending motor or segmental sensory afferent axon lengths in between stimulation and control groups (Supplemental Fig. S6). We conclude no difference in tissue-sparing or anatomical connections between the groups after SCI.

**Figure. 7.**
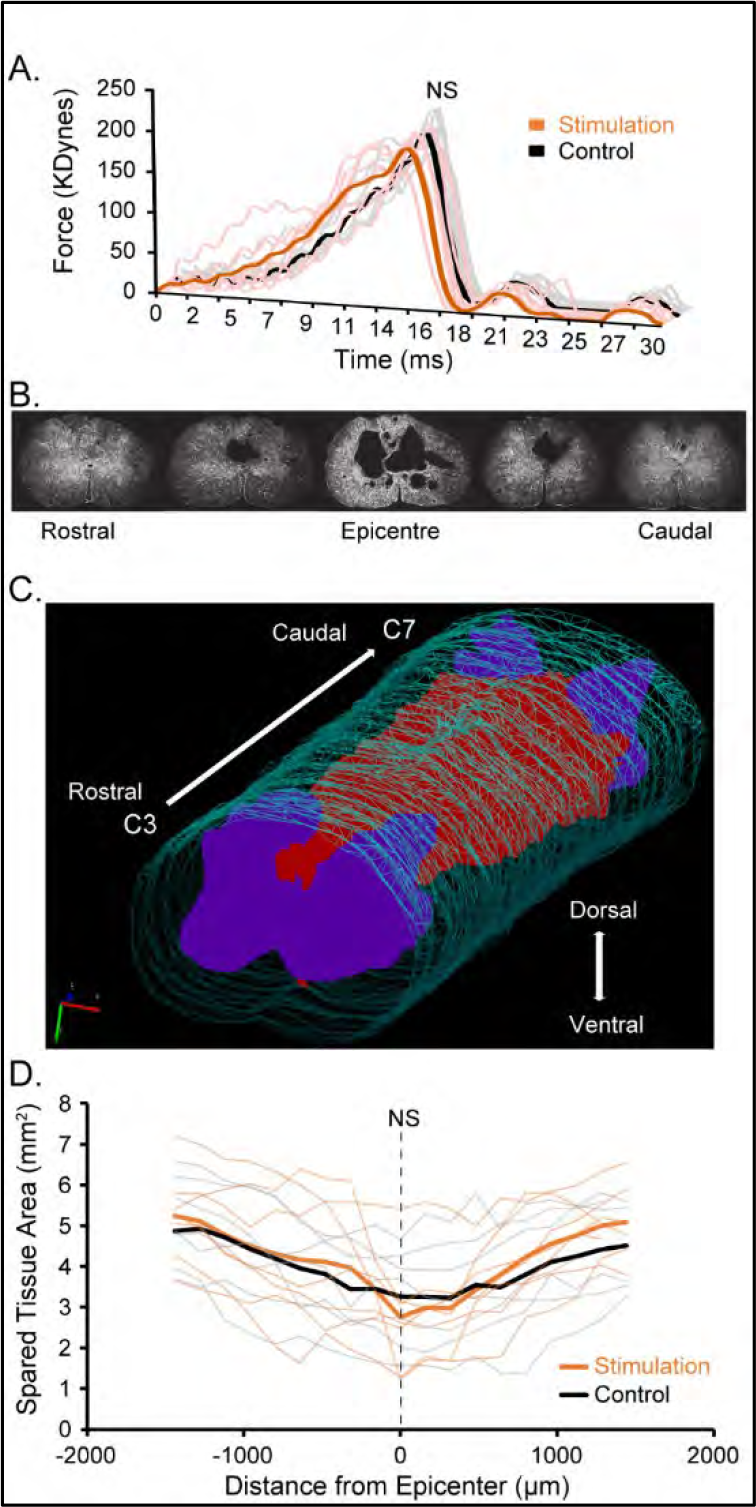
The injury force and the spared tissue did not differ between stimulation and control animals. (**A**) Force applied during cervical contusion (*n*=7 for stimulation, *n*=7 for control rats). (**B**) Representative spinal cord sections showing the extent of the lesion, scale bar: 1mm. (**C**) A rostro-caudal view of the lesion reconstruction. This 3D representation was created from cross sections through the site of spinal contusion. Lesion is shown in red, gray matter in purple, and the tissue border in green. Shows full degeneration of the main CST caudal to the site of injury, with partial sparing of the ventral pathways. (**D**) Spared tissue area through the lesioned spinal cord (*n*=7 for stimulation, *n*=7 for control rats).

## Discussion

Our main hypothesis proved true: repetitive pairing of motor cortex and spinal cord stimulation augmented spinal excitability and improved function in rats with SCI through spinal sensory and descending motor convergence. Paired stimulation produces effects on 3-time scales: immediate, hours, and weeks. The immediate effects (Fig. 1) do not represent plasticity; they operate only at the time that subthreshold spinal stimulation augments cortical MEPs. For paired stimulation to promote plasticity, it must be repeated (SCAP). Our optimized repetitive stimulation protocol produces robust augmentation that was still 50% greater than baseline at 2 hours after 30 minutes of pairing. Thus, some effects are short-lived, on the order of hours. However, when SCAP was delivered for 10 days, both function and physiology were improved as long as they were measured, to 50 days. Dexterity improved with no increase in pain (Fig. 4). Physiology changes included stronger MEPs with paired stimulation (Fig. 5) and improved H-reflex modulation (Fig. 6). These represent two critical functions of the sensorimotor system, to drive voluntary movement and to modulate segmental reflexes^39, 40^.

### SCAP depends on sensorimotor convergence in the spinal cord

Our model of SCAP as the convergence of descending sensory and segmental proprioceptors at the level of the spinal cord is supported by 3 lines of evidence. First, there is striking concordance between the time that motor potentials arrive in the spinal cord (9.17±0.55ms; Fig. 1A1) and the optimal latency for pairing for the immediate effects (10ms). Second, this immediate effect depends on the CST and proprioceptors, as demonstrated by abrogation of the effect with selective inactivation of each of these pathways (Fig. 2). Finally, pairing of motor cortex and spinal cord at a latency that has no immediate effect (100ms) also produces no plasticity when applied repetitively (Fig. 3D2 and supplementary Fig. 5).

We compared the effects of sensorimotor convergence at the level of the spinal cord with convergence in cortex, and the results challenge the most common clinical approach of targeting cortex. Paired afferent stimulation, which targets cortex, has been used to improve function in people, but the protocol does not work in all individuals, and the physiological effects are moderate in size^3, 41^. In contrast, convergence in the spinal cord had a much larger immediate and lasting effect in the rats. Several groups have applied paired stimulation of the descending (upper) motor system and segmental (lower) motor system^12–14, 42–45^. SCAP differs from these approaches by targeting descending motor and segmental afferents—sensorimotor integration rather than the motor only connections in the spinal cord.

### SCAP effective in rats with moderate cervical SCI

SCAP was at least as effective in rats with C4 SCI as uninjured rats, indicating that sparse descending motor circuits mediate the plasticity effect, along with the intact sensory system below injury. The C4 contusion injury obliterates the gray matter and leaves just a thin rim of white matter, like human injury^46, 47^. Interestingly, we observed sparing of <1% of the CST in rats with this injury^30^. This suggests that other descending motor pathways may be involved, as they are in recovery after human SCI^48^. Targeting the reticulospinal tract, which is slightly better preserved by experimental SCI in rats, results in improvement of motor function, both in rodents^23, 49^ and in humans^11, 50^.

There is a striking concordance of physiological changes induced by SCAP at the cortical and spinal cord levels. This is likely explained by changes in responses in the spinal cord, possibly in the common target of descending motor and segmental afferent connections. These results contribute to our working model that convergent inputs to spinal neurons cause the synapses or the cellular targets themselves to undergo lasting change. Synchrony of input is critical, since paired stimulation with an inappropriate time difference between motor cortex and spinal cord stimulation was ineffective in producing plasticity.

### Translation

This study provides important considerations for the application of paired stimulation to enhance sensorimotor function in people. The interactions between descending motor pathways and large-diameter afferents are largely conserved between rats and humans^51, 52^, but the direct connections between motor cortex and spinal motoneurons in humans may alter these interactions. In addition, the scale of the two species is very different, and it will be important to determine the proper timing of paired stimulation in people. Finally, it will be important to consider the differences in the location of tracts within the spinal cord and the pattern of injury to the spinal cord between human injury and experimental SCI in rats.

## Abbreviations

AAV: Adeno-Associated Virus
ADM: Abductor Digiti Minimi
CDP: cord dorsum potential
CNO: Clozapine-N-Oxide
CST: Corticospinal Tract
DREADD: Designer Receptor Exclusively Activated by Designer Drug
DRG: Dorsal Root Ganglion
IBB: Irvine, Beatties, and Bresnahan scale
MEP: Motor Evoked Potential
RGS: Rat Grimace Scale;
SCAP: Spinal Cord Associative Plasticity;
SCI: Spinal Cord Injury.

## Acknowledgements

We thank James McIntosh for helpful discussions about the statistical methods. We thank Tong Chun Wen for help with image acquisition. We thank Aldo Sandoval-Garcia and Walter Voit for making the spinal electrodes.

## Supplementary material

Supplementary material is available online.

## Supplementary material

### Materials and Methods

#### Descriptions of animals and surgical care

##### Animals

Fifty-five adult female Sprague-Dawley rats (Charles River, bodyweight of 275±25 grams) were included in this study. Out of these 31 rats were uninjured and 24 rats received spinal contusion. To test the proper timing of brain and spinal cord stimulation, a group of 9 healthy rats were tested on the immediate effect of adding subthreshold spinal cord stimulation to suprathreshold cortical stimulation (Fig. 1B), of these, 5 were used to test the effects of subthreshold stimulation both cortical and spinal (Fig. 1C). To understand which neural circuits mediate these effects, 8 rats had inactivation of neural circuits. Three rats were used for CST inactivation (Fig. 2A1-A4), two for sensory afferent inactivation (Fig. 2B1-B3), and Three female Long-Evans transgenic rats (250-275g), expressing Cre recombinase under the rat parvalbumin promoter {LE-TG(Pvalb-iCre)2Ottc, rat genome database (RGD) ID #10412329, RRRC line #773, NIDA, USA}, Pvalb-iCre rats were used for proprioceptor inactivation experiments (Fig. 2C1-C3). To test the effects of repetitive brain and spinal cord stimulation (SCAP), 14 healthy rats were used for the development and testing of SCAP (Figs. 3). To determine whether the neural circuits spared by SCI could support SCAP, 16 rats were subjected to a moderate cervical (C4) contusion. Two injured rats were initially used to test the short-term efficacy of SCAP. To test the long-term efficacy of SCAP to restore physiology and function, a group of 14 rats were randomized to real or sham (n=7, each) SCAP repeated over 10 days. Of the rats with a spinal contusion, 8 died immediately after SCI.

##### Surgery and Care

For all surgical procedures, rats were anesthetized via intraperitoneal injection of a mixture of ketamine (90mg/kg) and xylazine (10mg/kg body-weight). Local intradermal application of bupivacaine (0.25-0.5%, 0.1-0.2mL) was done before making skin incision and buprenorphine (0.05mg/kg body-weight, subcutaneous) was administered before and after the survival surgery and also after H-reflex testing to alleviate pain. Anesthesia levels were monitored by respiration and heart rate and responses to foot pinch. Animals were placed on heating pads (PhysioSuite, Kent Scientific Corporation, Northwest CT, USA) to maintain a body temperature of 37.5°C as measured continuously by a rectal probe (PhysioSuite). The animals were housed in individual cages with free access to food and water on a 12/12-hour light/dark cycle. All experimental procedures were in full compliance with the approved Institutional Animal Care and Use Committee Protocol of Columbia University.

#### Cortical, EMG and spinal electrodes

In a single procedure, motor cortex screw electrodes and EMG wires were placed and the head connectors brought to the common head cap. For motor cortex electrodes, rats were anesthetized and head-fixed in a stereotaxic frame (David Kopf Instruments, Tujunga, CA, USA). The skull was exposed and placement holes made using a Jobber Drill (number 60, P1 Technologies, Roanoke, VA, USA) without disturbing the dura mater. Stainless steel screw electrodes (1.19mm diameter with a flat tip; Plastics One, Torrington, CT, USA) were implanted over the forelimb area of the motor cortex in one hemisphere at two locations as described in our previous work^1^. The screw electrodes were attached in advance to a head connector (Plastics One) that was secured with skull screws and dental acrylic. For electromyogram (EMG) recordings, we inserted supple stainless steel braided wire (diameter=0.014mm; #AS 634, Cooner Wire Co., Chatsworth, CA, USA) electrodes into the biceps muscle (for MEPs) and *adductor digiti minimi* (#AS 631, for H-reflex). For the biceps, the other end was tunneled subcutaneously and attached in advance to a head connector, mounted on the skull, and also secured with skull screws and dental acrylic (Lang Dental Manufacturing Co., Inc., Wheeling, IL, USA).

One week after a cortical and EMG electrode implant, the rats went through another surgery to implant spinal stimulation electrodes, as described previously^2, 3^. Briefly, the head and the cervical spine were fixed in a stereotactic frame. The T1 spinous process was clamped to stabilize the spine. The C4 spinal cord was exposed by laminectomy. Array electrodes were placed in the dorsal epidural space in the midline of the spinal cord over C5-C6. Muscles and skin were sutured in layers. The array head connector was mounted on the skull and also secured with skull screws and dental acrylic.

#### MEP measurements

All the MEP measurements were done in awake behaving rats in their home cages. Before the start of testing, electrodes were connected and the animal was left in the cage for 10-15 minutes to get habituated. We stimulated the motor cortex and measured MEPs from contralateral biceps muscle. Each motor cortex was stimulated independently and contralateral EMGs acquired from the biceps. A similar primary motor cortex and spinal cord stimulation paradigm were used as described by Mishra et al., 2017^1^. Briefly, a train of three biphasic square wave pulses (each pulse of 0.2ms for each polarity; interstimulus interval of 3ms; Fig. 1B) was used for motor cortex stimulation. Multichannel EMG signals were acquired before and after the repetitive stimulation at 5,000Hz sampling rate, with a CED (CED Micro 1401, Cambridge Electronic Design Ltd, Cambridge, UK) data acquisition system and a recording software (Signal 5.08, CED Ltd). The signals were amplified at a gain of x1000 and band pass filtered 1–1000Hz with an integrated AC differential amplifier system (A-M Systems, Model 1700, Sequim, WA, USA). Motor cortex and spinal stimulation were performed using an isolated pulse stimulator (A-M Systems, Model 2100). MEPs were extracted, processed, and quantified using customized Motometrics analysis^4^.

For the motor cortex stimulation we first measured the cortical threshold, defined as the stimulation intensity needed to evoke a short latency MEP in >50% of trials, by stimulating cortex at increasing intensities. We then measured the MEP at 110% of the cortical threshold, which we call cortical MEPs. The same method was then used to determine the spinal threshold and to measure MEPs at 110% of spinal threshold, which we call spinal MEPs. For the paired stimulation, we stimulated the motor cortex at 110% of cortical threshold and the spinal cord was stimulated with 90% spinal threshold intensity. To quantify MEPs, the EMGs were rectified and then averaged across 20 trials. The area-under-the-curve (AUC) was calculated for MEPs and % change from baseline was quantified. To determine the most effective latency, we stimulated the spinal cord at various time intervals from 30ms before motor cortex stimulation to 30ms after motor cortex stimulation. Latencies were tested in a randomized sequence, with a minimum of 2 minutes between each test. For pairing at each latencies we first determined the cortical and spinal thresholds before the repetitive paired stimulation and 110% of cortical thresholds was used for baseline MEP recordings, which was motor cortex stimulation alone. For pairing we used 110% of cortical threshold and 90% of spinal threshold intensities. At each latencies we measured spinal threshold and pairing was done at 90% of the threshold using baseline cortical threshold. For both Subthreshold stimulation, intensities for both cortical and spinal stimulations were set to 90% of their threshold values. First, as control recordings, only-spinal and only-cortical stimulations were applied to record the MEP AUC during these subthreshold stimulations. Then, paired stimulations at 90% of thresholds with latencies between ±30ms were applied, and MEPs were recorded.

#### Corticospinal tract and spinal afferent inactivation

##### Corticospinal tract (CST) inactivation

The CST was selectively inactivated using a Designer Receptor Exclusively Activated by Designer Drug (DREADD), during paired stimulation. To target the CST selectively, a Cre-dependent adeno-associated virus (AAV1) encoding the DREADD was injected into the motor cortex and another encoding Cre-recombinase was injected into the cervical spinal cord. Thus, only doubly infected neurons express the DREADD, which can be inactivated by the administration of its ligand, clozapine-N-oxide (CNO)^5^. Through a craniotomy, twelve injections of 200nl of AAV1-hSyn-DIO-hM4D(Gi)-mCherry {2×10^12^ vg/ml, pAAV-hSyn-DIO-hM4D(Gi)-mCherry was a gift from Bryan Roth (Addgene viral prep # 44362-AAV1 ; http://n2t.net/addgene:44362 ; RRID:Addgene_44362)^6^ were made 1.5mm from the pial surface into the forelimb area of the left motor cortex at the following forelimb motor cortex coordinates^7^: AP 0.5 ML 1.5; AP 0.5, ML 2.5; AP 0.5, ML 3.5, AP 1.5, ML 1.5; AP 1.5, ML 2.5; AP1.5, ML 3.5; AP 2.5, ML 1.5; AP 2.5, ML 2.5; AP 2.5, ML 3.5; AP 3.5, ML 2.0, AP 3.5, ML 3.0; AP 4.5, ML2.5. AAV retrograde; AAV2 pmSyn1-EBFP-Cre (5.5 x 10^12^ vg/ml, AAV pmSyn1-EBFP-Cre was a gift from Hongkui Zeng; Addgene plasmid # 51507; http://n2t.net/addgene:51507 ; RRID:Addgene_51507^8^ was injected into the right side of the cervical spinal cord. Eleven injections of 200nl of AAV2 were made in C5 and C6, 0.8mm lateral from the midline at the depth of 1.2mm from the surface of the spinal cord, with 0.6mm intervals between injections.

One week after AAV injections, animals were trained on the vermicelli handling task^9, 10^. After reaching a stable baseline performance, the number of forepaw manipulations was measured before, during, and after inactivation. The number of adjustments of each forepaw was counted while the animal was eating uncooked pasta before and at 30, 60, and 120 minutes after intraperitoneal injection of 2.4mg/kg body weight of clozapine N-oxide (CNO). One week after the behavioral test, we tested the effects of inactivation on the immediate effects of paired brain and spinal cord stimulation. Under deep anesthesia a cranial window was created to expose the forelimb motor cortex, and silver ball electrodes were placed over two locations: 1.0mm anterior, and 2.0mm lateral; and 3.0mm anterior and 4.0mm lateral to bregma. After that we exposed the C4-C6 vertebrae, and laminectomy was done at C5-C6 level to expose dorsal surface of spinal cord and a pair of silver ball electrodes were placed epidurally on the midline of C5-C6 cervical spinal cord. A pair of EMG wires were inserted in the biceps muscle bilaterally to record EMG. Systemic administration of CNO typically causes maximum inactivation of DREADD expressing neurons within 30 minutes, and CNO typically washes out within two hours^11^. This allows testing of physiology and behavior before, during, and after reversible inactivation of specific neural circuits. The MEP of each biceps was tested by independent stimulation of the contralateral motor cortex. Measurements were taken every 10 minutes for 2 hours. The cortical threshold changed during this time, so paired stimulation was always compared with cortex-only stimulation at the corresponding time point.

##### Afferent inactivation

The necessity of afferents for the immediate effects of paired stimulation was tested using two inactivation strategies. For the first strategy, large diameter dorsal root ganglion (DRG) neurons were inactivated. Large DRG neurons mediate touch, proprioception, and muscle length/tension^12^. The specificity of large-diameter DRG neurons was provided by the tropism of the AAV5^13^. One microliter of AAV5-hSyn-hM4D(Gi)-mCherry (2×10^13^ vg/ml, pAAV-hSyn-hM4D(Gi)-mCherry was a gift from Bryan Roth (Addgene viral prep # 50475-AAV5; http://n2t.net/addgene:50475 ; RRID:Addgene_50475) was injected into each DRG at C5, C6, and C7 on one side of the spinal cord after partial laminectomy. At the time of virus injection, stainless steel screw electrodes were placed over each motor cortex, spinal array on the epidural surfaces of C5-C6 cervical spinal cord, and EMG wires in the biceps muscle bilaterally. Similar to CST inactivation, four to five weeks after the virus injection, large-diameter afferents were inactivated by an intraperitoneal injection of 2.4mg/kg body weight of CNO and MEPs measured over 2 hours. For the second strategy, we targeted the proprioceptive afferents for inactivation, since these mediate the effects of spinal cord stimulation^14, 15^. We used a genetic strategy to target DRG neurons that express parvalbumin, which are proprioceptors^16, 17^. In transgenic rats that express Cre recombinase under the control of parvalbumin promoter PV-Cre rats (LE-Tg(Pvalb-iCre)2Ottc, Optogenetics and Transgenic Technology Core, NIDA, MD, USA obtained via Rat Research Resource Centre RRRC#773; the genotype was confirmed prior to virus injections by PCR using the primers, Pvalb forward: 5’-CCG TTC AGC TTG AAG GTG TC-3’, iCre R12: 5’-CAC AGT CAG CAG GTT GGA GA-3’) we injected a Cre-dependent DREADD (AAV5-hSyn-DIO-hM4D(Gi)-mCherry (2×10^13^ vg/ml, pAAV-hSyn-DIO-hM4D(Gi)-mCherry was a gift from Bryan Roth; Addgene viral prep # 44362-AAV5; http://n2t.net/addgene:44362 ; RRID:Addgene_44362^6^ into the C5, C6, and C7 DRGs as described above. Inactivation with CNO and physiology measurement were also identical. In these rats cortical and EMG electrodes were implanted one week before DRG injections, and the spinal array was implanted just after the injection using the methods described above.

##### Inactivation controls

For CNO only control, immediate effects of paired stimulation was recorded from right and left forelimb biceps before CNO injection in a normal rat (no virus injection). After CNO injection immediate effects of paired stimulation was observed every 10 minute till 2 hours. For saline only control in two PV-Cre rats immediate effects of paired stimulation was recorded from both right and left forelimb biceps before normal saline (sterile injectable 0.9%NaCl, vehicle for CNO) injection. After saline injection immediate effects of paired stimulation effects was recorded every 10 minute till 2 hours. The cortical and spinal MEPs, as well as cortical and Spinal excitability were calculated as described below. The excitability was expressed as 1/ threshold, data was normalized to baseline values before injection.

#### SCAP protocol and optimization

##### SCAP protocol

We tested brain and spinal cord responses at baseline and then after the period of repetitive pairing (Fig. 3A1-A3), as we did in Mishra et al., 2017^1^ but in awake rats with implanted electrodes. This differs from the immediate effects of stimulation (Fig. 1A3). Before modulation, we measured the stimulation threshold for the motor cortex to produce a MEP in 50% of trials and also the spinal threshold. We then stimulated with 110% of threshold to determine cortical and spinal MEPs, using the area under the curve. At baseline, cortical and spinal MEPs, as well as cortical and spinal thresholds were measured singly and independently (Fig. 3A1). For repetitive pairing (Fig. 3A2), motor cortex stimulation was performed at 110% of the threshold for evoking a cortical MEP followed 10ms later by spinal cord stimulation performed at 90% of the threshold for generating a spinal MEP. After repetitive pairing (Fig. 3A3), the same measures taken at baseline were taken immediately after pairing and every 20 min thereafter up to 120 min. We quantified excitability as 1/threshold and expressed it as % change from baseline. Add description of each session threshold testing and quantification of excitability. Here we determined the cortical and spinal thresholds before the repetitive paired stimulation and 110% of cortical and spinal thresholds were used for baseline MEP recordings. For SCAP we used 110% of cortical threshold and 90% of spinal threshold intensities. After SCAP, each time points till 120 minutes we measured both spinal and cortical thresholds to calculate excitability (1/threshold) and MEP testing was done similar to baseline testing by stimulating cortex and spinal cord at 110% of threshold that was determined before SCAP.

##### SCAP optimization

In order to maximize the efficacy of the SCAP protocol (5-minute SCAP, Fig 3B1)^1^, we varied four parameters in order to produce the largest physiological responses: frequency, number of stimuli, number of trains, and time between trains. In two rats implanted with cortical and spinal electrodes, we first varied frequency by testing 10 stimuli delivered at 5Hz, 2Hz, 1Hz, 0.5Hz and 0.2Hz (Supplementary Fig. 3A). We have tested whether repeated stimuli suppressed the resulting MEP. For the other parameters, we examined the short term SCAP effects with the average percent increase in cortical MEPs, cortical and spinal excitability over the 120 minutes after the SCAP protocol (Fig. 3B2). Spinal excitability was used since this was less variable than spinal MEPs, and out of concern that eliciting spinal MEPs might induce plasticity. In addition, we intended to keep the total stimulation period to 30 minutes, a period that could be used for stimulation in people. For these trials, we used 5 Hz pairing and varied the number of stimuli by comparing 150, 75 & 38 pulses (Supplementary Fig. 3B, C & D1-D3). For subsequent trials we used 20 trains and varied the time between trains, either 1 or 5 minutes (Supplementary Fig. 3B, C & E1-E3). Further, for subsequent trials we used 150 stimuli, which we term a train, and varied the number of trains, either 10 or 20 (Supplementary Fig. 3B, C & F1-F3). The optimized 30-minute SCAP protocol is shown in Fig. 3B2. A train of 150 stimuli at 5Hz frequency (30 seconds) is followed by a 60-second gap. The train and gap is then repeated 20 times, delivering 3000 stimuli pairs in 30 minutes.

#### C4 spinal contusion and SCAP after SCI

##### Cervical spinal cord contusion

The C4 spinal cord contusion surgery was performed as previously described^18^. Briefly, the C3 to C5 vertebrae were exposed by skin incision and muscle separation on the midline. Laminectomy of C4 was performed, taking care to ensure the dura remained intact. Following laminectomy, the rat was placed on the Infinite Horizon (IH) impactor stabilization platform (IH-0400, Precision Systems and Instrumentation, LLC), and the C3 and C5 spinous processes were clamped. A fine adjustment was made to the vertebral clamps to ensure that the exposed C4 spinal cord surface was level. The moderate contusion injury was made with a 3.5mm impactor tip raised to three complete turns ∼2mm from the dorsal surface and was dropped with a preset 200kdynes force on to the spinal cord with no dwell time. The force curve of the impactor was saved so that the rats in the Stim and Control groups could be compared. After injury, the rat was removed from the platform and the spinal electrode array was implanted as described above. Rats were individually housed in separate cages provided with ALPHA-dri dust-free animal bedding (Shepherd Specialty Papers, Watertown, TN) and received intense care after SCI. This included placing their cage on a heating pad for 12h, administering buprenorphine (4 doses in total, 0.05mg/kg) every 8–12h, and Baytril (5mg/kg, Norbrook, Henry Schein, Melville, NY) daily for 5 days. For nutrition, Ringer’s Lactate solution (10ml, subcutaneous) was administered daily for 5–10 days and Diet boost Gel (ClearH2O, Portland, ME) was placed within reach until rats regained the ability to reach the food and water dispensers. Body weight, food and water intake was monitored daily. Bladders were palpated every 12 hours; none of the rats showed bladder dysfunction.

##### SCAP in SCI rats

We designed a preclinical efficacy trial, as shown in Fig. 4A. Ten days of an optimized SCAP protocol was delivered in the subacute period, 10 days after injury^18, 19^. The daily stimulation duration (30 minutes) and number of sessions (10) were designed to be clinically viable. The primary outcome measure was the dexterity (IBB) score, and secondary outcomes included changes in MEPs, H-reflex modulation, and a pain measure (RGS). The study was powered based on the effect size we observe of a different neuromodulation strategy^18^. A-priori power analysis suggested we need 7 rats in each group to achieve a power of 0.75. However, the physiological effects of SCAP were more robust than the previous paradigm, suggesting a stronger behavioral effect. The post-hoc power with a total of 14 rats was 0.85 using the same parameters as the initial power calculation.

#### Behavior testing

##### Food manipulation task (IBB)

To assess the rats ability to manipulate, we used the Irvine, Beatties, and Bresnahan scale^20, 21^, as we have done before^18^. All rats were placed in a clear plastic cylinder on top of a rotating carousel so that they could be rotated in the direction of the video camera. The rats were habituated to the testing environment for 10 days, 15 minute per day. For testing, rats were video recorded while eating both donut-shaped (Froot Loops, Kellogg’s NA Co.) and sphere-shaped (Reese’s Puffs, General Mills Sales Inc.) cereal. Experimenters blinded to treatment recorded the rats eating the cereals before SCI and weekly from 4 to 7 weeks after SCI. Analysis of the recorded session was done offline by a blinded observer. The scores for the two different kinds of cereal and the two forepaws were averaged. A schematics showing characteristic features of rat forelimbs during cereal manipulation task is presented in Fig. 4B1. Images of a representative stimulation rat were generated from frames of live videos with the image tracing tool of Adobe Illustrator software.

##### Horizontal ladder walking task

The horizontal ladder walking task measured paw placement on the rungs, which were irregularly spaced^19, 22, 23^. After cortical electrode implantation, we trained rats 20 min daily for 10 days to ensure rats walked across the ladder without distractions. We motivated rats to cross the ladder with puffs from a can of compressed air and by offering a cotton swab dipped into 20% sucrose after every successful trial. The pattern of the irregularly spaced rungs was altered every 6 trials and the walking direction was changed after 12 trials. We video recorded the behavior at 50 frames-per-second. Experimenters blinded to the paired stimulation treatment analyzed the trials frame-by-frame to quantify the step quality. The start was defined when the rat placed all 4 limbs on the rungs and the end when it reached the last rung of the ladder. The criteria were used to establish when to begin and end scoring and also to measure the time taken to walk across the ladder. The steps were scored as a good step, overstep, under-step, or a missed step based on the placement of the forepaw on the rung; for the main analysis, this was divided only into good steps or errors. We excluded one rat that did not meet the baseline average of 15% error rate

##### Rat grimace scale (RGS)

This scale is used to quantify pain using the rats facial characteristics^24^, the way that is commonly used to assess pain in people. All rats were kept on a rotating carousel as in the case of the food manipulation task, and the facial features were recorded with a camera by a blinded experimenter at the times shown in Fig. 4. The facial features were analyzed offline by a blinded scorer.

#### H-reflex testing

To track H-reflex modulation, we measured the H-reflex at baseline, one week after SCI, just after 10 days of stimulation and every two weeks post-stimulation (Fig.4A). The M-wave is evoked by the excitation of motor axons. The H-reflex is the electrophysiological equivalent of the spinal stretch reflex; Ia proprioceptive afferents synapse on and activate motor neurons in the spinal cord (Fig. 6A). Rats were lightly anesthetized with 60 mg/kg ketamine. A pair of braided wire electrodes were inserted across the medial plantar side of the wrist to stimulate the ulnar nerve (via a constant current isolated pulse stimulator, single biphasic pulse, width 100µs, A-M Systems). Two recording electrodes were inserted into the abductor digiti minimi to record EMGs. The threshold was determined as the lowest stimulation intensity that elicited an H-wave response in at least 75% of the recordings^25^. First, we tested responses to increasing stimulus intensities up to 2X threshold, which activates low threshold afferent fibers. Next, we tested the frequency-dependent depression of the H-wave^25–27^. This is also commonly referred to as rate-dependent depression. We stimulated every 10s with paired stimuli at inter-stimulus intervals from 10s to 0.01s. The H-wave area under the curve (AUC) of the test stimulus was normalized to the H-wave AUC of the conditioning stimulus. 25 paired stimuli per frequency were averaged and plotted as a frequency-depression curve (Fig. 6C & D).

#### Histological analysis

To determine the severity of SCI, we compared the parameter (at the time of SCI) and severity of injury between those with and without therapeutic stimulation. One week later, rats were deeply anesthetized and perfused, first with ∼400ml 0.1M phosphate-buffered saline (PBS) containing 5,000 UPS units/ml of heparin, and then with 600 ml 4% paraformaldehyde. Spinal cord (C1-T2) were dissected and post-fixed in 4% paraformaldehyde for 48h. The tissue was then transferred to 30% sucrose at 4°C until it was cut in the coronal section at 40μm thickness on a cryostat. Tissue sections from C3 to C5 were processed for lesion reconstruction. Spinal cord sections were collected serially and directly onto slides to preserve tissue integrity. The tissue sections dried overnight and were coverslipped the next day. The outline of the lesion, grey matter, and white matter were traced using dark-field microscopy and Stereo-investigator software (MBF Bioscience, Williston, Vermont, USA). A blinded experimenter used the Planimetry tool to reconstruct sections around the lesion epicenter to determine the lesion and total tissue section area around the lesion epicenter. The spared tissue area was computed by subtracting the lesion tissue area from the total tissue area. To compare the extent of injury in the two groups, spared tissue area data was quantified every 200 microns across the whole area^18^.

##### Axon Quantification

After collecting the behavior and physiology data, a BDA tracer was injected into one-half of the primary motor cortex. The BDA injections were done either one week after AAV1 (in DRG) injections or one week before CTb (in nerve). For BDA injections the dental acrylic head cap was removed with an electric drill, and the electrode screws were extracted with a screwdriver. After the electrodes were taken out, a 3×3mm craniotomy window was created above the right side forelimb area motor cortex in order to inject the tracer. 10% biotinylated dextran amine (BDA; 10,000mw, Molecular Probes, Eugene, OR) was injected at depth of 1.5mm at the following 7 sites, 500nl each site (mm rostral to bregma, mm lateral to bregma): (2.0, 2.5), (1.0 2.5), (0.5 3.0), (1.5 3.0), (2.5 3.0), (2.0 3.5), (1.0 3.5). After the injections, a thin layer of GelFoam (Pfizer, NY) was placed over the dura, and the wound was closed.

For axon quantification, BDA histology was performed on sections above (C3) and below (C7) the lesion site. Briefly, sections from spinal cord levels C3 and C7 were rinsed with 0.1M PBS, then incubated for 2h in Alexa Fluor 594 streptavidin (1:500 in PBS; Invitrogen #11227). After 2h of incubation sections were rinsed with PBS and sections were mounted on charged glass slides using Fluoroshield (Sigma-Aldrich # F6182) for preserving fluorescence of tissue, then sections were cover-slipped. Slides were stored at 4^0^C till visualization. For each rat, data were collected from above and below the lesion site as described. Both the ipsilateral and the contralateral sides of the C7 sections were hand traced using Stereoinvestigator (MBF Bioscience, Williston, Vermont, USA). To correct for differences in the efficacy of the tracer, axon lengths were normalized by the efficiency of tract tracing^19, 22, 28^. The optical fractionator probe in Stereoinvestigator was used to determine an estimate of the number of labeled axons in the contralateral dorsal column at C3. A correction factor was calculated by dividing each rat’s averaged dorsal column axon number by the average number of axons in the dorsal column. Six rats were excluded from the BDA quantification analysis (three from each groups) because of too little BDA staining to be quantified due to the ineffective batch of BDA.

For transganglionic tracing of the central projections of sensory axons, the median nerves was exposed under sterile conditions, and 1–2μL of a 1% (wt/vol) solution of CTb (List Laboratories) was injected using a Hamilton syringe 7 day before the rats were perfused^29^. For visualization of CTb, sections were stained with goat anti-CTb (1:2000; List Laboratories) and anti-goat conjugated to FITC (1:500; Millipore)^30^. We did not observed any CTb labeled axons in the spinal cord, so in next cohort of rats we injected AAV1 for sensory axon quantification. For sensory axon quantifications, AAV1 was injected in C5-C7 DRG as described above (7.74 x10^13^; pAAV-hSyn-EGFP was a gift from Bryan Roth (Addgene viral prep # 50465-AAV1; http://n2t.net/addgene: 50465; RRID: Addgene_50465) and sensory afferent axons were quantified below the lesion segments with the Stereoinvestigator (MBF Bioscience, Williston, Vermont, USA). At spinal cord level C7, the space balls probe was used to estimate the axon length in the grey matter ipsilateral to the Virus injection. To correct for differences in tracer efficiency, axon length was normalized by computing the number axons in the ipsilateral dorsal column using the optical fractionator probe and a correction factor was calculated in the same way as for the motor axons.

#### Randomization and blinding procedures

Rats were pseudorandomized into real (Stim) and sham (Control) groups with each cohort of up to 8 rats having rats assigned to each group. Rats were randomized immediately after SCI surgery, and experiments were carried out with blinded measures. The experimenter who randomized rats into different groups and performed paired stimulation treatment, but was not involved in behavior tasks training, testing or scoring process or in histology. Experimenters who performed behavior training, testing, and scoring and histology did not participate in randomization, electrophysiology testing, or paired stimulation. Control rats were connected to the stimulator, but no stimulation was delivered. Since all rats had both cortical and spinal electrodes implanted, there was no way to distinguish by appearance whether rats received stimulation.

### Statistical analyses

All of the comparisons between stimulation conditions were done with paired/unpaired t-tests with Bonferroni correction or One-way ANOVA with Bonferroni corrections and Cohen’s d was used effect size. To determine the latency for paired stimulation (Fig.1), we compared MEPs for pairing at different latencies to the baseline (only motor cortex stimulation at 110% of threshold). For the sub-threshold timing, the baseline MEPs were normalized and compared to the pairing at different latencies. For the inactivation experiments (Figs. 2), the immediate effects of the pairing on the targeted side at baseline was compared to the different times after CNO injection. For SCAP (Fig. 3), we used only descriptive statistics as percentage change in MEP. Independent cortex and spinal MEPs were compared to each time after SCAP. For the preclinical randomized trial, the Stim and Control groups were compared by an independent t-test with Bonferroni correction. The immediate effects of pairing was compared between the groups at several time points (Fig. 5). H-reflex modulation was compared only at the shortest (0.01s) inter-stimulus-interval (Fig. 6). The dexterity and pain scores were compared at the end of the trial (Fig. 4; day 49). To compare the injuries (Fig. 7), the peak force delivered to induce SCI (mean±SEM force at the time of impact was presented) and percentage spared tissue area at the epicenter between the groups was compared independent t-test. All the statistical analysis was performed with SPSS (SPSS Statistics for Windows, version 27; IBM Corp., Armonk, NY, USA). We tested the normality of the data using the Shapiro-Wilk test. All data are expressed as Mean±SEM.

## Results

### Motor axon quantification

For motor axon quantification, BDA histology was performed on sections above (C3) and below (C7) the lesion site. Some rats were excluded from the BDA quantification analysis because of too little BDA staining to be quantified (n=3 from each group) because the BDA dye was ineffective. In stimulated group total length of motor axons below the lesion were 3.1±1.44mm (*n*=4) while in control group it was 3.1±0.54mm (*n*=4; Supplementary Fig. S6A). We observed quantifiable labeling of BDA in 8 rats (4 in each group), in these rats, motor axons were quantified as described above. Three rats from each group were excluded in the motor axon quantification and analysis due to poor /no BDA staining that resulted due to the use of an ineffective batch of BDA.

### Sensory axon quantification

For transganglionic tracing of the central projections of sensory axons, first we tried CTb labelling and staining. We did not observed any CTb labeled axons in the spinal cord, so in the next cohort of rats, we injected AAV1 for sensory axon quantification. For the sensory axon quantifications, AAV1 was injected in C5-C7 DRG and sensory afferent axons were quantified below the lesion segments. The sensory axons were quantified only in the last cohort of animals comprising 2 rats in stimulation groups and 3 rats in the control group. The total sensory axon length below the lesion in the stimulation group was 76.4±26mm (*n*=2), and that in the control group was 94.9±52mm (*n*=3; Supplementary Fig. S6B). For sensory axon labeling, first, we used CTb in 9 rats. Out of these 9 rats, 2 animals (one from each group) died either after CTb injection or before perfusion. Spinal cord sections from 7 rats (stim, n=4; control, n=3) were looked for CTb immunostaining, but we only observed very poor/ no staining. So, we injected AAV1 in 5 rats (stim, n=2; control, n=3) and sensory axon quantifications were done in these rats.

## Supplementary Figure Legends

**Supplementary Figure 1.**
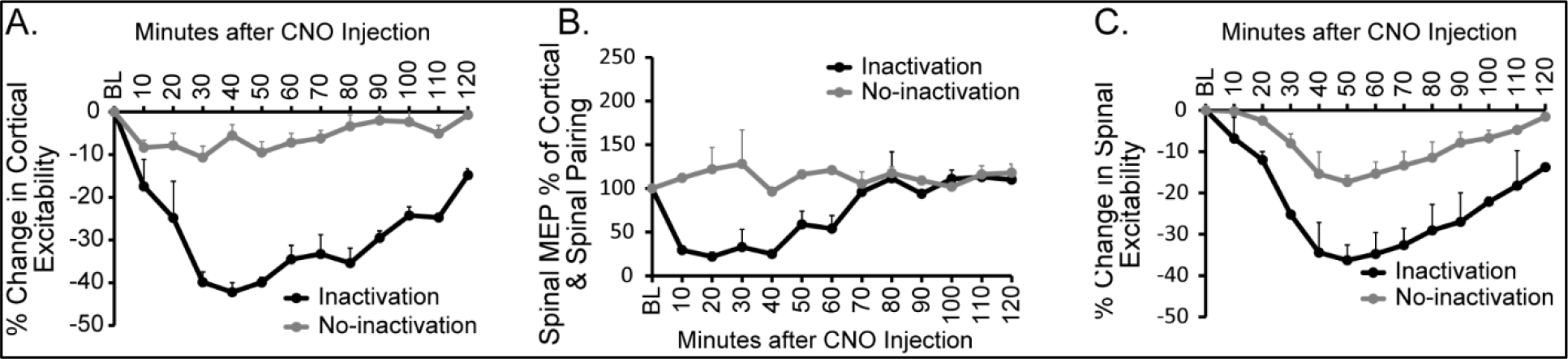
(A) Cortical excitability decreased after inactivation. Inactivation decreases the motor cortex excitability in PV-Cre rats, inactivation side (black line) compared to baseline (BL, *n* = 2); more than 40% suppression of motor cortex excitability at 40 minutes after CNO injection. While on the non-inactivated side less than 10% change in cortical excitability was observed. (**B)** Spinal MEP decreased after inactivation. Inactivation abrogates paired stimulation effects on spinal evoked MEPs in PV-Cre rats, inactivation side (black line) compared to baseline (BL, *n*=2); more than 70% suppression of paired stimulation effect at 20 minutes after CNO injection. While on the non-inactivated side no change in spinal MEPs observed. (**C)** Spinal excitability decreased after inactivation. Inactivation also decreases the spinal cord excitability in PV-Cre rats, inactivation side (black line) compared to baseline (BL, *n*=2); more than 30% suppression of spinal excitability at 40-50 minutes after CNO injection. While in the non-inactivated side less than 20% change in spinal excitability was observed. Error bars show SEM, excitability = 1/ threshold.

**Supplementary Figure 2.**
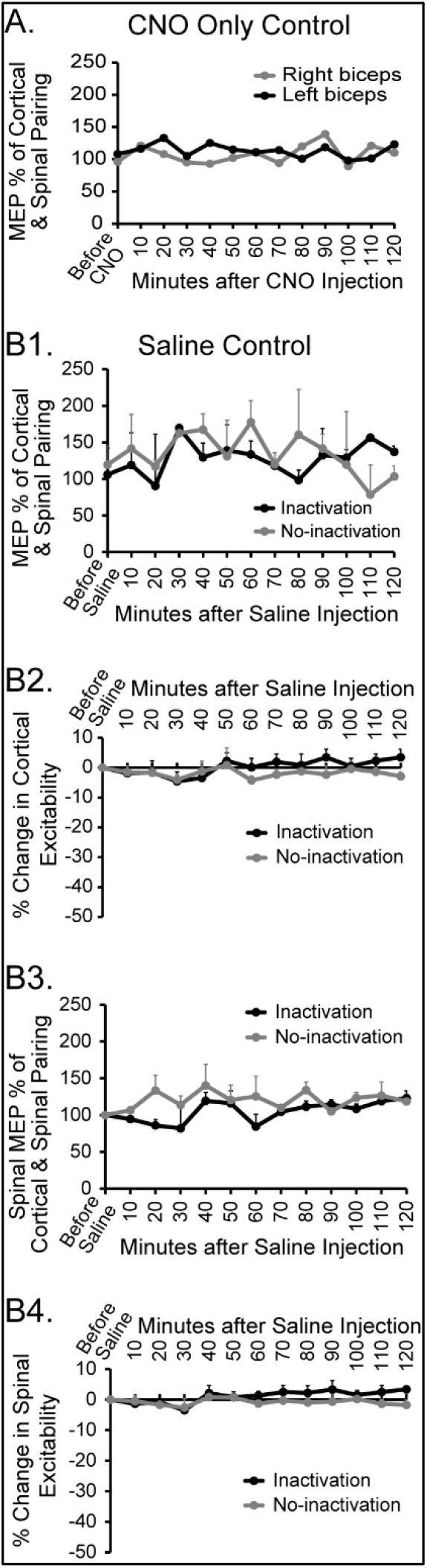
Inactivation Controls. **(A) CNO only control.** Immediate effects of paired stimulation were recorded from right (gray line) and left (black line) forelimb biceps before CNO injection in a normal rat (no virus injection). After CNO injection paired stimulation effects were recorded every 10 minutes for 2 hours. No change in paired stimulation effects on cortical MEPs was observed compared to before CNO injection. (**B1) Saline only control.** In two PV-Cre rats immediate effects of paired stimulation were recorded from right (gray line = no-inactivation side) and left (black line = inactivation side) forelimb biceps before normal saline (0.9%NaCl, vehicle for CNO) injection. After saline injection paired stimulation effects were recorded every 10 minutes for 2 hours. No change in paired stimulation effects on cortical MEPs was observation compared to before saline injection. (**B2)** Similarly cortical excitability was calculated before and after saline injection. No change in cortical excitability was observed compared to before saline injection. Similarly spinal MEPs (**B3**) and Spinal excitability (**B4**) also did not change after saline injection compared to before injection. Error bars show SEM, excitability = 1/ threshold.

**Supplementary Figure 3.**
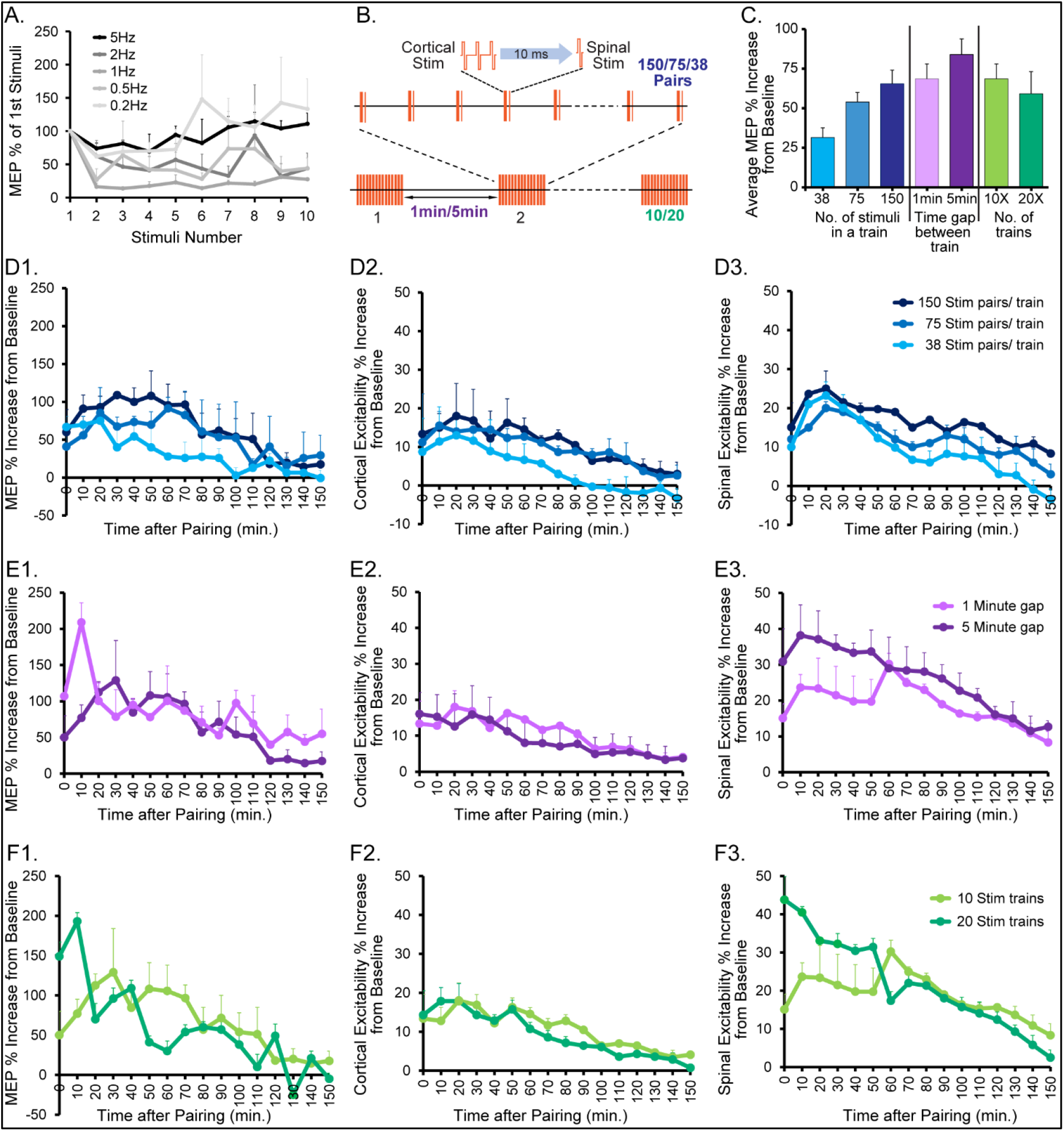
SCAP protocol optimization. **(A)** Effect of second stimulation over the first stimulation for different frequencies. (**B)** Protocol to test the effects of different numbers of paired pulse trains, time intervals between the pulse trains and repeated trains. **(C)** Effects of number of stimuli in a train (150 stimuli are optimum), time interval between trains (1 min was as good as 5 min) and number of trains (10x and 20x are similar) on average MEPs over 120 minutes of testing. (**D1 – D3)** Using the 5Hz frequency, we tested for the length of the train, 150 stimuli were more effective than the others tested. (**E1 – E3) For** the time interval between two bursts, we observed that the 5-minute interval was slightly better than 1 minute, but we selected 1 minute to minimize the total duration of stimulation. (**F1 – F2)** Finally, we tested the total number of repetitions of trains. We observed that 20 trains of stimulation had a more durable effect than 10 trains with a similar overall magnitude. Error bars show SEM, excitability = 1/ threshold.

**Supplementary Figure 4.**
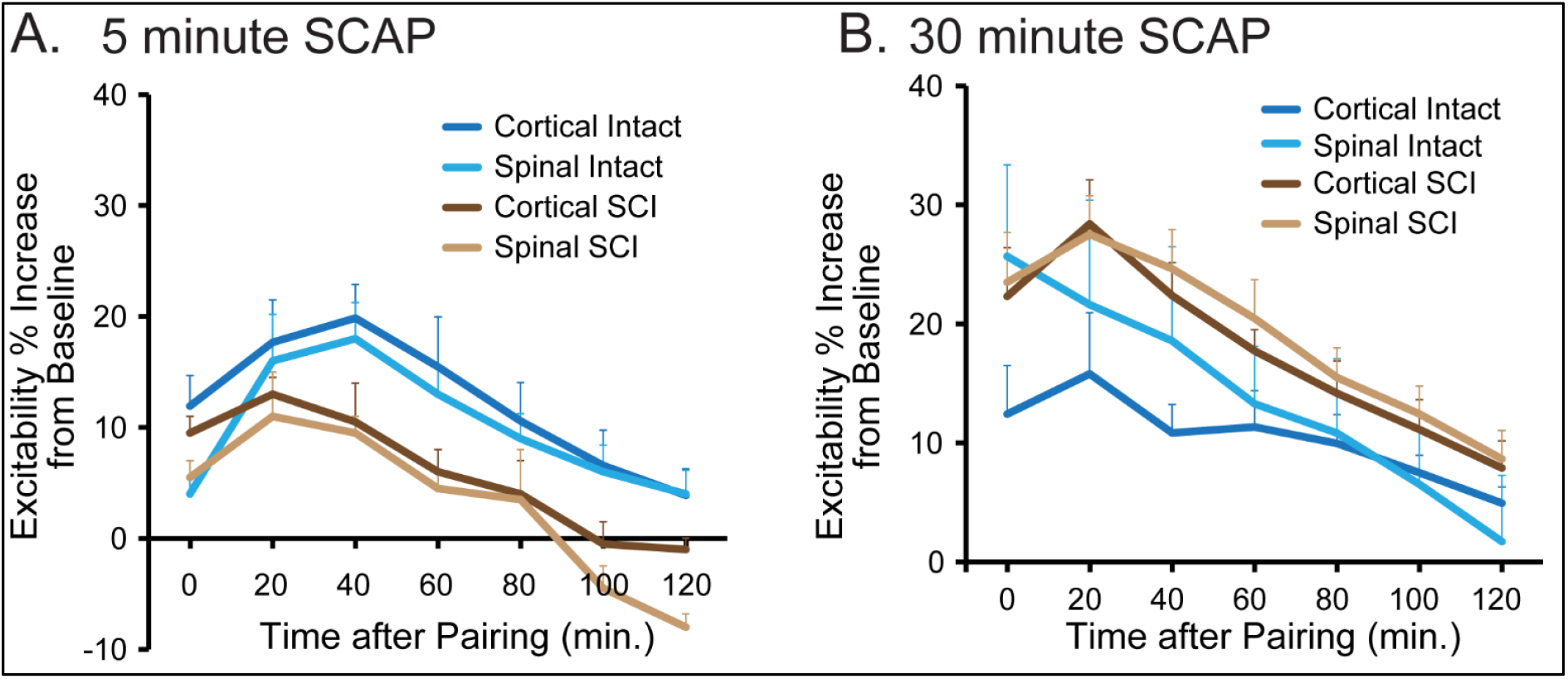
SCAP causes lasting changes in cortical and spinal cord excitability in uninjured rats and rats with SCI. (**A)** Cortical and spinal excitability after 5-minute SCAP in intact rats (*n*=6) as well as in rats with C4 SCI (*n*=2). In both SCI and intact groups of rats after 5-minutes of repetitive pairing, both cortical and spinal excitability was augmented for more than 1 hour and this augmentation persisted even after SCI, then returned to the baseline. (**B)** Cortical and spinal excitability after 30-minute SCAP in intact rats (*n*=4) and in SCI rats (*n*=7). In both groups of rats after 30-minutes of repetitive pairing, both cortical and spinal excitability was augmented for more than 2 hours and this augmentation persisted even after SCI, similarly the spinal MEP and excitability was also increased for 2 hours after pairing. Error bars show SEM, excitability = 1/ threshold.

**Supplementary Figure 5.**
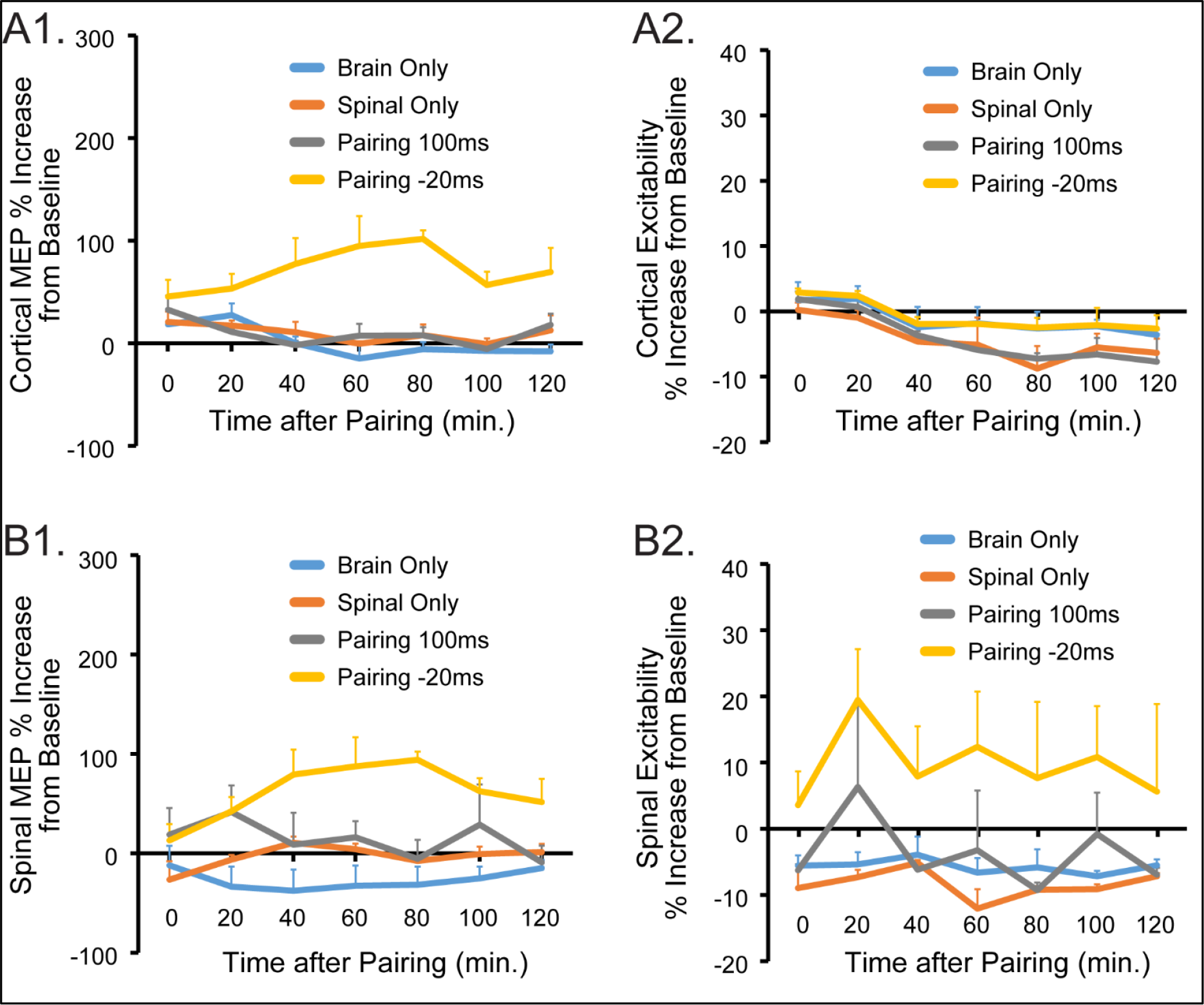
SCAP controls. (A1) Effect of 30-minute SCAP in controlled conditions. Baseline cortical MEP, spinal MEP and thresholds were measured before 30-minue repetitive pairing. Cortical and spinal stimulation at 110% of the cortical and spinal thresholds. MEPs recorded from the biceps. Threshold was determined by adjusting stimulation intensity to provoking a short-latency EMG in >50% of trials. For only brain stimulation, motor cortex was stimulated for 30-minutes repetitively at 100% of cortical threshold, after 30 minutes of stimulation baseline measurements were done at each time-point for 2 hours. Similarly for spinal only, spinal cord was stimulated for 30-minutes at 90% spinal threshold and after that baseline measurements were done at each time-point for 2 hours. For pairing at 100ms, motor cortex and spinal cord stimulation were paired repetitively for 30-minutes. Here, first motor cortex was stimulated (at 110% of cortical threshold) then 100ms latter spinal cord was stimulated (at 90% of spinal threshold). After 30 minutes of stimulation, baseline measurements were done at each time-point for 2 hours. To pair motor cortex and spinal cord stimulation at −20ms, first spinal cord was stimulated at 90% of spinal threshold, then 20ms latter motor cortex was stimulated at 110% of cortical threshold and repeated for 30-minutes. After 30 minutes of repetitive stimulation, baseline measurements were done at each time-point for 2 hours. (**A1)** There was a slight increase in cortical MEPs just after repetitive pairing in all the conditions and it came back to baseline quickly except for pairing at −20ms, here it was at an elevated level for 2hours. (**A2)** The cortical excitability was not changed as compared to baseline values in all the four different conditions. (**B1)** The spinal MEPs also follows similar trends to cortical MEPs but with more variability. (**B2)** Spinal excitability also follows the same trends to cortical excitability except for pairing at −20ms, here there was lots of variability between the rats. Error bars show SEM, excitability = 1/ threshold.

**Supplementary Figure 6.**
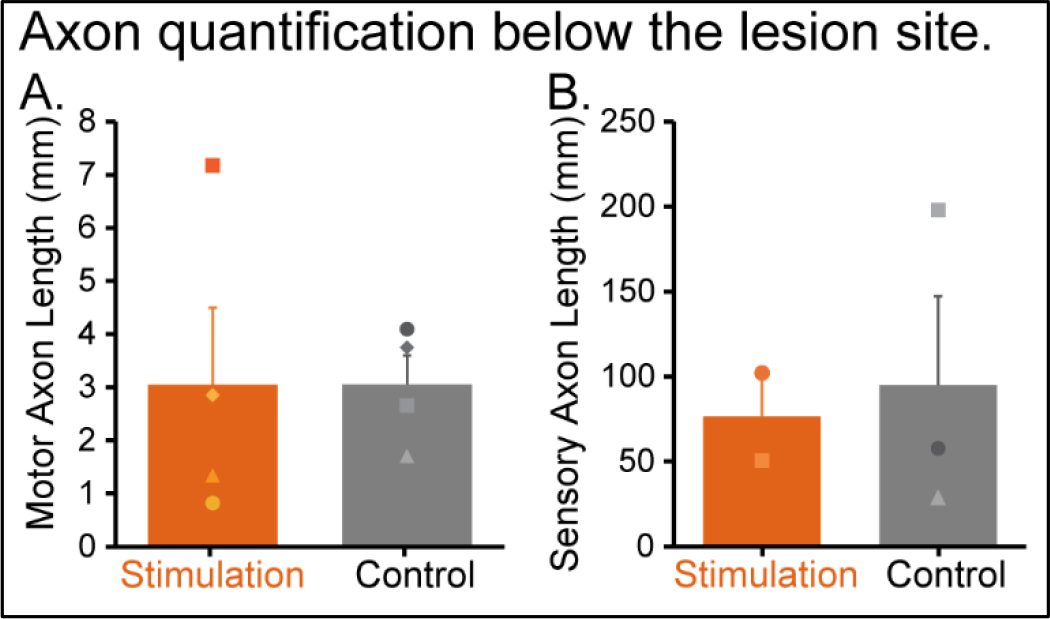
Total motor and sensory axon lengths below the lesion in both stimulation and control animals. (**A**) In the stimulated group total length of motor axons below the lesion were 3.1±1.44mm (n=4) while in the control group it was 3.1± 0.54mm (n=4). (**B**) The total sensory axon length below the lesion in the stimulation group were 76.4±26mm (n=2), and that in the control group was 94.9± 52mm (n=3). Error bars show SEM.

**Supplementary table 1:**
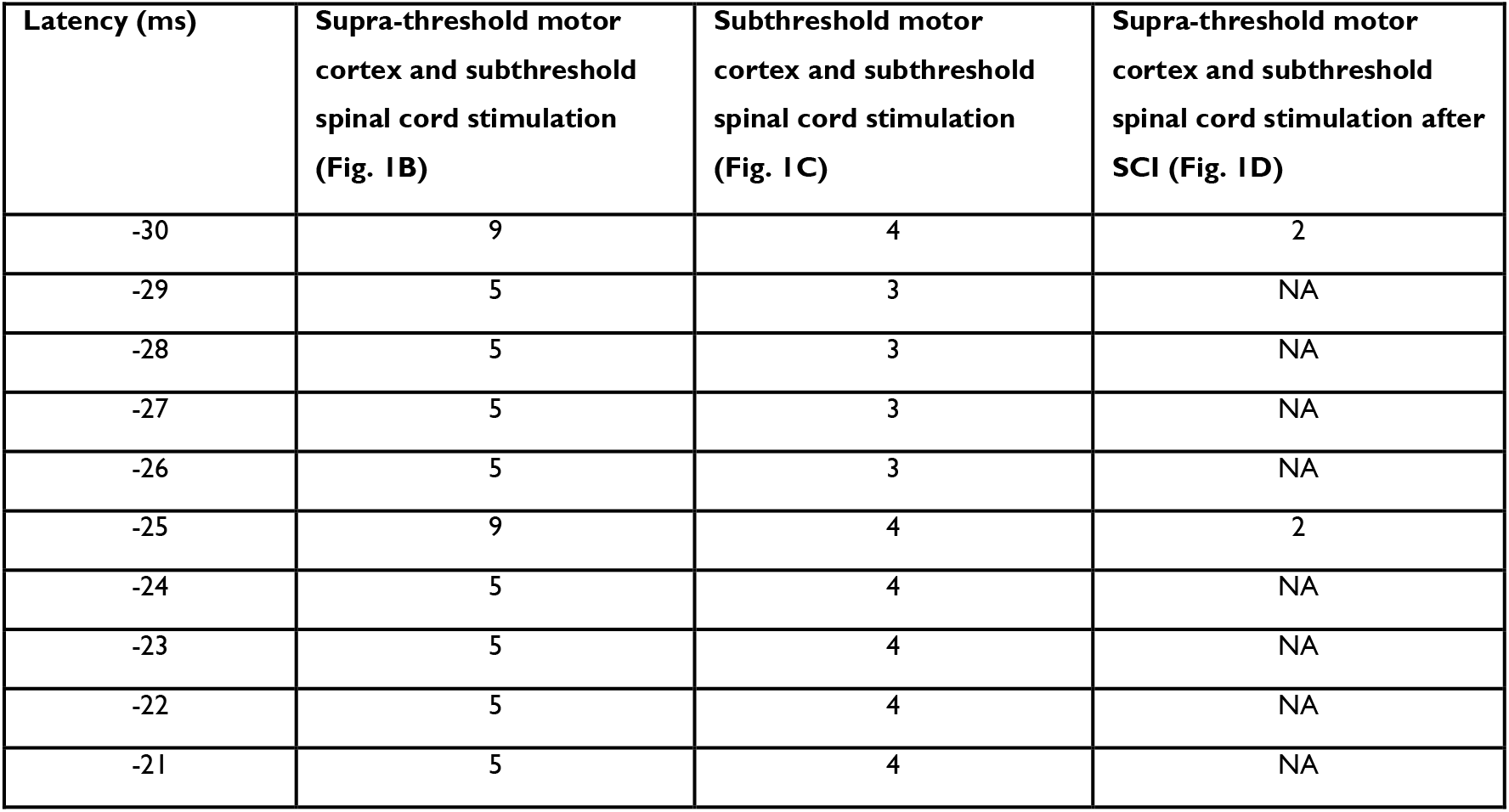
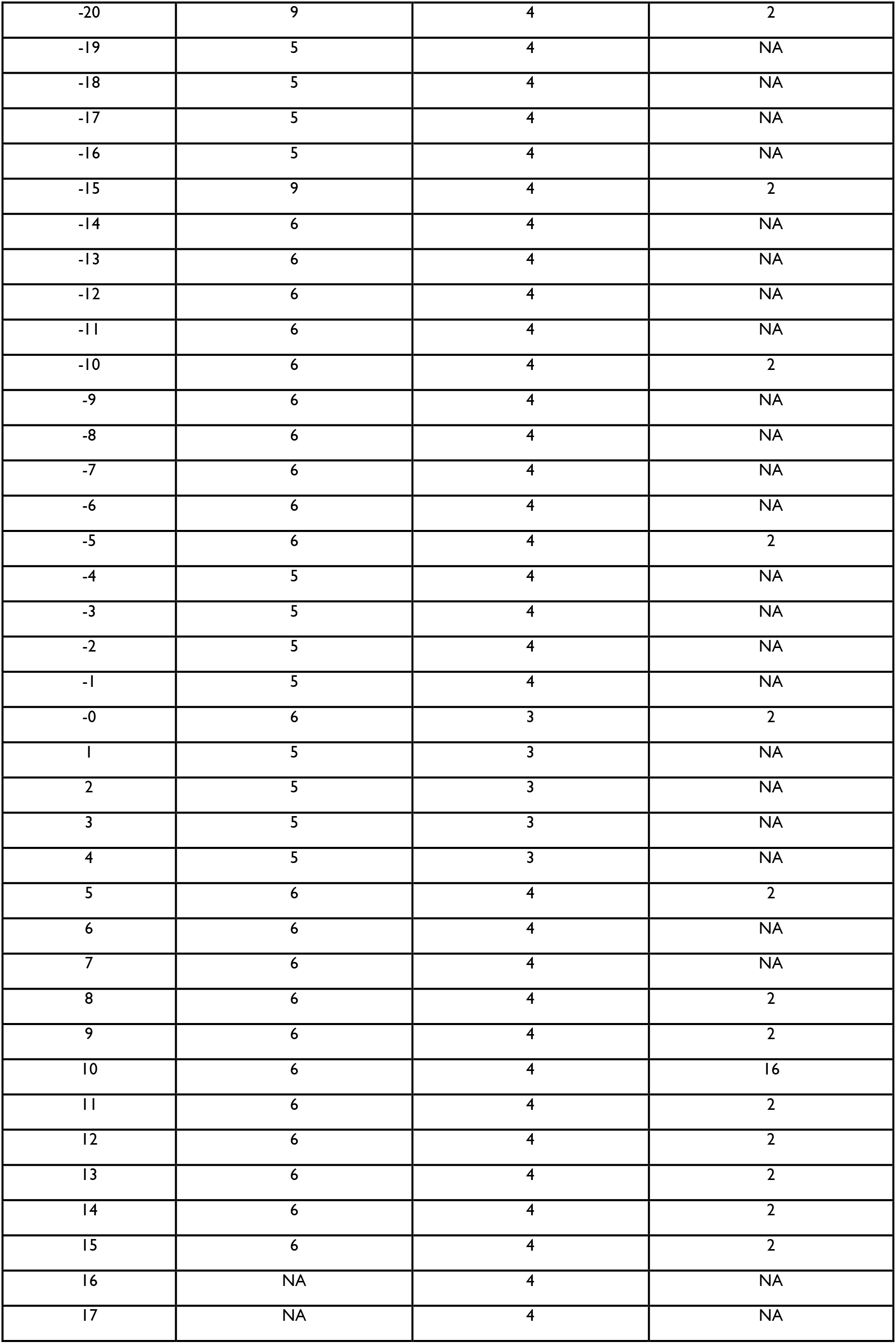
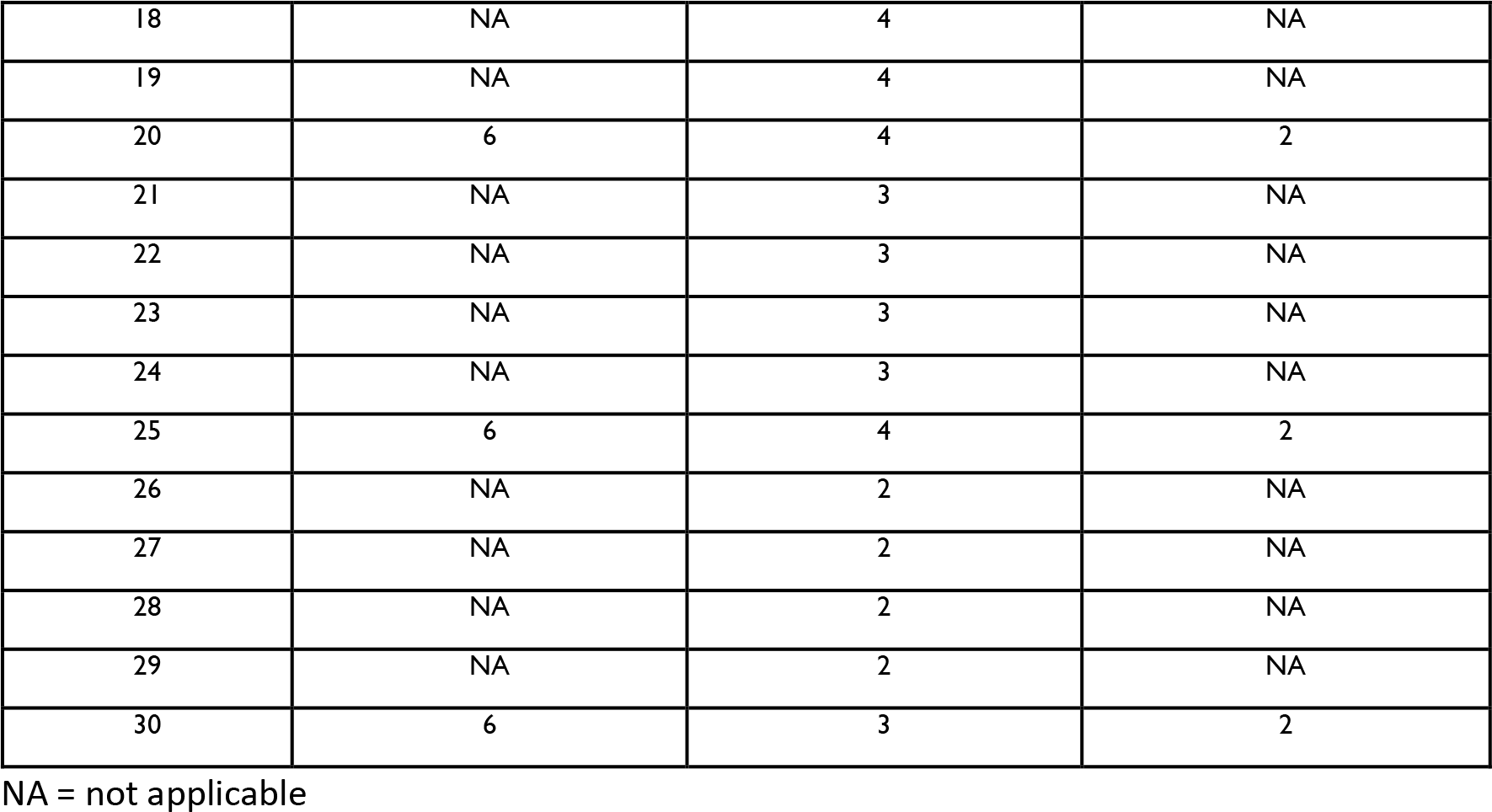
Number of animals (n) used in the immediate effect of motor cortex and spinal cord stimulation latency testing.

## References

1. Asan AS, McIntosh JR, Carmel JB. Targeting Sensory and Motor Integration for Recovery of Movement after CNS Injury. Front Neurosci. 2022 Jan 21;15:791824. doi: 10.3389/fnins.2021.791824

2. Stefan K, Kunesch E, Cohen LG, Benecke R, Classen J. Induction of plasticity in the human motor cortex by paired associative stimulation. Brain. 2000;123 Pt 3:572–584. doi:10.1093/brain/123.3.572

3. Suppa A, Quartarone A, Siebner H, et al. The associative brain at work: Evidence from paired associative stimulation studies in humans. Clin Neurophysiol. 2017;128(11):2140–2164. doi:10.1016/j.clinph.2017.08.003

4. Ling YT, Alam M, Zheng Y-P. Spinal Cord Injury: Lessons about Neuroplasticity from Paired Associative Stimulation. Neuroscientist. 2020;26(3):266–277. doi:10.1177/1073858419895461

5. Wagner FB, Mignardot J-B, Le Goff-Mignardot CG, et al. Targeted neurotechnology restores walking in humans with spinal cord injury. Nature. 2018;563(7729):65–71. doi:10.1038/s41586-018-0649-2

6. Angeli CA, Boakye M, Morton RA, et al. Recovery of Over-Ground Walking after Chronic Motor Complete Spinal Cord Injury. N Engl J Med. 2018;379(13):1244–1250. doi:10.1056/NEJMoa1803588

7. Gill ML, Grahn PJ, Calvert JS, et al. Neuromodulation of lumbosacral spinal networks enables independent stepping after complete paraplegia. Nat Med. 2018;24(11):1677–1682. doi:10.1038/s41591-018-0175-7

8. Nscisc. National Spinal Cord Injury Statistical Center (Facts and Figures at a Glance). 2016.

9. Anderson KD. Targeting recovery: priorities of the spinal cord-injured population. J Neurotrauma. 2004;21(10):1371–1383. doi:10.1089/neu.2004.21.1371

10. Bunday KL, Perez MA. Motor recovery after spinal cord injury enhanced by strengthening corticospinal synaptic transmission. Curr Biol. 2012;22(24):2355–2361. doi:10.1016/j.cub.2012.10.046

11. Bunday KL, Urbin MA, Perez MA. Potentiating paired corticospinal-motoneuronal plasticity after spinal cord injury. Brain Stimulat. 2018;11(5):1083–1092. doi:10.1016/j.brs.2018.05.006

12. Jo HJ, Perez MA. Corticospinal-motor neuronal plasticity promotes exercise-mediated recovery in humans with spinal cord injury. Brain. May 2020. doi:10.1093/brain/awaa052

13. Nishimura Y, Perlmutter SI, Eaton RW, Fetz EE. Spike-timing-dependent plasticity in primate corticospinal connections induced during free behavior. Neuron. 2013;80(5):1301–1309. doi:10.1016/j.neuron.2013.08.028

14. Nishimura Y, Perlmutter SI, Fetz EE. Restoration of upper limb movement via artificial corticospinal and musculospinal connections in a monkey with spinal cord injury. Front Neural Circuits. 2013;7:57. doi:10.3389/fncir.2013.00057

15. Mishra AM, Pal A, Gupta D, Carmel JB. Paired motor cortex and cervical epidural electrical stimulation timed to converge in the spinal cord promotes lasting increases in motor responses. J Physiol (Lond). 2017;595(22):6953–6968. doi:10.1113/JP274663

16. Garcia-Sandoval A, Pal A, Mishra AM, et al. Chronic softening spinal cord stimulation arrays. J Neural Eng. 2018;15(4):045002. doi:10.1088/1741-2552/aab90d

17. Garcia-Sandoval A, Guerrero E, Hosseini SM, et al. Stable softening bioelectronics: A paradigm for chronically viable ester-free neural interfaces such as spinal cord stimulation implants. Biomaterials. 2021;277:121073. doi:10.1016/j.biomaterials.2021.121073

18. Park HG, Carmel JB. Selective manipulation of neural circuits. Neurotherapeutics. 2016;13(2):311–324. doi:10.1007/s13311-016-0425-7

19. Wahl AS, Omlor W, Rubio JC, et al. Neuronal repair. Asynchronous therapy restores motor control by rewiring of the rat corticospinal tract after stroke. Science. 2014;344(6189):1250–1255. doi:10.1126/science.1253050

20. Allred RP, Adkins DL, Woodlee MT, et al. The vermicelli handling test: a simple quantitative measure of dexterous forepaw function in rats. J Neurosci Methods. 2008;170(2):229–244. doi:10.1016/j.jneumeth.2008.01.015

21. Wen T-C, Lall S, Pagnotta C, et al. Plasticity in one hemisphere, control from two: adaptation in descending motor pathways after unilateral corticospinal injury in neonatal rats. Front Neural Circuits. 2018;12:28. doi:10.3389/fncir.2018.00028

22. Giuffrida R, Rustioni A. Dorsal root ganglion neurons projecting to the dorsal column nuclei of rats. J Comp Neurol. 1992;316(2):206–220. doi:10.1002/cne.903160206

23. Vulchanova L, Schuster DJ, Belur LR, et al. Differential adeno-associated virus mediated gene transfer to sensory neurons following intrathecal delivery by direct lumbar puncture. Mol Pain. 2010;6:31. doi:10.1186/1744-8069-6-31

24. Takeoka A, Vollenweider I, Courtine G, Arber S. Muscle spindle feedback directs locomotor recovery and circuit reorganization after spinal cord injury. Cell. 2014;159(7):1626–1639. doi:10.1016/j.cell.2014.11.019

25. Takeoka A. Proprioception: Bottom-up directive for motor recovery after spinal cord injury. Neurosci Res. 2020;154:1–8. doi:10.1016/j.neures.2019.07.005

26. Asboth L, Friedli L, Beauparlant J, et al. Cortico-reticulo-spinal circuit reorganization enables functional recovery after severe spinal cord contusion. Nat Neurosci. 2018;21(4):576–588. doi:10.1038/s41593-018-0093-5

27. Ichikawa H, Deguchi T, Nakago T, Jacobowitz DM, Sugimoto T. Parvalbumin, calretinin and carbonic anhydrase in the trigeminal and spinal primary neurons of the rat. Brain Res. 1994;655(1-2):241–245. doi:10.1016/0006-8993(94)91620-9

28. Walters MC, Sonner MJ, Myers JH, Ladle DR. Calcium imaging of parvalbumin neurons in the dorsal root ganglia. eNeuro. 2019;6(4). doi:10.1523/ENEURO.0349-18.2019

29. Krashes MJ, Koda S, Ye C, et al. Rapid, reversible activation of AgRP’’ neurons drives feeding behavior in mice. J Clin Invest. 2011;121(4):1424–1428. doi:10.1172/JCI46229

30. Yang Q, Ramamurthy A, Lall S, et al. Independent replication of motor cortex and cervical spinal cord electrical stimulation to promote forelimb motor function after spinal cord injury in rats. Exp Neurol. 2019;320:112962. doi:10.1016/j.expneurol.2019.112962

31. Carmel JB, Berrol LJ, Brus-Ramer M, Martin JH. Chronic electrical stimulation of the intact corticospinal system after unilateral injury restores skilled locomotor control and promotes spinal axon outgrowth. J Neurosci. 2010;30(32):10918–10926. doi:10.1523/JNEUROSCI.1435-10.2010

32. Irvine K-A, Ferguson AR, Mitchell KD, et al. The irvine, beatties, and bresnahan (IBB) forelimb recovery scale: an assessment of reliability and validity. Front Neurol. 2014;5:116. doi:10.3389/fneur.2014.00116

33. Irvine K-A, Ferguson AR, Mitchell KD, Beattie SB, Beattie MS, Bresnahan JC. A novel method for assessing proximal and distal forelimb function in the rat: the Irvine, Beatties and Bresnahan (IBB) forelimb scale. J Vis Exp. 2010;(46). doi:10.3791/2246

34. Carmel JB, Kimura H, Martin JH. Electrical stimulation of motor cortex in the uninjured hemisphere after chronic unilateral injury promotes recovery of skilled locomotion through ipsilateral control. J Neurosci. 2014;34(2):462–466. doi:10.1523/JNEUROSCI.3315-3.2014

35. Carmel JB, Martin JH. Motor cortex electrical stimulation augments sprouting of the corticospinal tract and promotes recovery of motor function. Front Integr Neurosci. 2014;8:51. doi:10.3389/fnint.2014.00051

36. Metz GA, Whishaw IQ. Th. e ladder rung walking task: a scoring system and its practical application. J Vis Exp. 2009;(28). doi:10.3791/1204

37. Sotocinal SG, Sorge RE, Zaloum A, et al. The Rat Grimace Scale: a partially automated method for quantifying pain in the laboratory rat via facial expressions. Mol Pain. 2011;7:55. doi:10.1186/1744-8069-7-55

38. Tan AM, Chakrabarty S, Kimura H, Martin JH. Selective corticospinal tract injury in the rat induces primary afferent fiber sprouting in the spinal cord and hyperreflexia. J Neurosci. 2012;32(37):12896–12908. doi:10.1523/JNEUROSCI.6451-11.2012

39. Fitzgerald PB. An update on the clinical use of repetitive transcranial magnetic stimulation in the treatment of depression. J Affect Disord. 2020;276:90–103. doi:10.1016/j.jad.2020.06.067

40. Porter R, Lemon R. Corticospinal function and voluntary movement. Oxford University Press; 1993. doi:10.1093/acprof:oso/9780198523758.001.0001

41. Wolpaw JR. The negotiated equilibrium model of spinal cord function. J Physiol (Lond). 2018;596(16):3469–3491. doi:10.1113/JP275532

42. Carson RG, Kennedy NC. Modulation of human corticospinal excitability by paired associative stimulation. Front Hum Neurosci. 2013 Dec 3;7:823. doi: 10.3389/fnhum.2013.00823

43. Taylor JL, Martin PG. Voluntary motor output is altered by spike-timing-dependent changes in the human corticospinal pathway. J Neurosci. 2009;29(37):11708–11716. doi:10.1523/JNEUROSCI.2217-09.2009

44. Harel NY, Carmel JB. Paired stimulation to promote lasting augmentation of corticospinal circuits. Neural Plast. 2016;2016:7043767. doi:10.1155/2016/7043767

45. Chandrasekaran S, Nanivadekar AC, McKernan G, et al. Sensory restoration by epidural stimulation of the lateral spinal cord in upper-limb amputees. Elife. 2020;9. doi:10.7554/eLife.54349

46. Bunge RP, Puckett WR, Becerra JL, Marcillo A, Quencer RM. Observations on the pathology of human spinal cord injury. A review and classification of 22 new cases with details from a case of chronic cord compression with extensive focal demyelination. Adv Neurol. 1993;59:75–89

47. Kakulas BA, Kaelan C. The neuropathological foundations for the restorative neurology of spinal cord injury. Clin Neurol Neurosurg. 2015;129 Suppl 1:S1–7. doi:10.1016/j.clineuro.2015.01.012

48. Baker SN, Perez MA. Reticulospinal Contributions to Gross Hand Function after Human Spinal Cord Injury. J Neurosci. 2017;37(40):9778–9784. doi:10.1523/JNEUROSCI.3368-16.2017

49. Moritz CT. Now is the critical time for engineered neuroplasticity. Neurotherapeutics. 2018;15(3):628–634. doi:10.1007/s13311-018-0637-0

50. Sangari S, Lundell H, Kirshblum S, Perez MA. Residual descending motor pathways influence spasticity after spinal cord injury. Ann Neurol. 2019;86(1):28–41. doi:10.1002/ana.25505

51. Curfs MH, Gribnau AA, Dederen PJ, Bergervoet-Vernooij IW. Induction of c-fos expression in cervical spinal interneurons after kainate stimulation of the motor cortex in the rat. Brain Res. 1996;725(1):88–94.

52. Pierrot-Deseilligny E, Burke D. The circuitry of the human spinal cord: its role in motor control and movement disorders. Cambridge: Cambridge University Press; 2005. doi:10.1017/CBO9780511545047

## Reference

1. Mishra AM, Pal A, Gupta D, Carmel JB. Paired motor cortex and cervical epidural electrical stimulation timed to converge in the spinal cord promotes lasting increases in motor responses. J Physiol (Lond). 2017;595(22):6953–6968. doi:10.1113/JP274663

2. Garcia-Sandoval A, Pal A, Mishra AM, et al. Chronic softening spinal cord stimulation arrays. J Neural Eng. 2018;15(4):045002. doi:10.1088/1741-2552/aab90d

3. Garcia-Sandoval A, Guerrero E, Hosseini SM, et al. Stable softening bioelectronics: A paradigm for chronically viable ester-free neural interfaces such as spinal cord stimulation implants. Biomaterials. 2021;277:121073. doi:10.1016/j.biomaterials.2021.121073

4. Ratnadurai Giridharan S, Gupta D, Pal A, Mishra AM, Hill NJ, Carmel JB. Motometrics: A toolbox for annotation and efficient analysis of motor evoked potentials. Front Neuroinformatics. 2019;13:8. doi:10.3389/fninf.2019.00008

5. Park HG, Carmel JB. Selective manipulation of neural circuits. Neurotherapeutics. 2016;13(2):311–324. doi:10.1007/s13311-016-0425-7

6. Krashes MJ, Koda S, Ye C, et al. Rapid, reversible activation of AgRP’’ neurons drives feeding behavior in mice. J Clin Invest. 2011;121(4):1424–1428. doi:10.1172/JCI46229

7. Brus-Ramer M, Carmel JB, Martin JH. Motor cortex bilateral motor representation depends on subcortical and interhemispheric interactions. J Neurosci. 2009;29(19):6196–6206. doi:10.1523/JNEUROSCI.5852-08.2009

8. Madisen L, Garner AR, Shimaoka D, et al. Transgenic mice for intersectional targeting of neural sensors and effectors with high specificity and performance. Neuron. 2015;85(5):942–958. doi:10.1016/j.neuron.2015.02.022

9. Allred RP, Adkins DL, Woodlee MT, et al. The vermicelli handling test: a simple quantitative measure of dexterous forepaw function in rats. J Neurosci Methods. 2008;170(2):229–244. doi:10.1016/j.jneumeth.2008.01.015

10. Wen T-C, Lall S, Pagnotta C, et al. Plasticity in one hemisphere, control from two: adaptation in descending motor pathways after unilateral corticospinal injury in neonatal rats. Front Neural Circuits. 2018;12:28. doi:10.3389/fncir.2018.00028

11. Wahl AS, Omlor W, Rubio JC, et al. Neuronal repair. Asynchronous therapy restores motor control by rewiring of the rat corticospinal tract after stroke. Science. 2014;344 (6189):1250–1255. doi:10.1126/science.1253050

12. Giuffrida R, Rustioni A. Dorsal root ganglion neurons projecting to the dorsal column nuclei of rats. J Comp Neurol. 1992;316(2):206–220. doi:10.1002/cne.903160206

13. Vulchanova L, Schuster DJ, Belur LR, et al. Differential adeno-associated virus mediated gene transfer to sensory neurons following intrathecal delivery by direct lumbar puncture. Mol Pain. 2010;6:31. doi:10.1186/1744-8069-6-31

14. Takeoka A, Vollenweider I, Courtine G, Arber S. Muscle spindle feedback directs locomotor recovery and circuit reorganization after spinal cord injury. Cell. 2014;159(7):1626–1639. doi:10.1016/j.cell.2014.11.019

15. Asboth L, Friedli L, Beauparlant J, et al. Cortico-reticulo-spinal circuit reorganization enables functional recovery after severe spinal cord contusion. Nat Neurosci. 2018;21(4):576–588. doi:10.1038/s41593-018-0093-5

16. Ichikawa H, Deguchi T, Nakago T, Jacobowitz DM, Sugimoto T. Parvalbumin, calretinin and carbonic anhydrase in the trigeminal and spinal primary neurons of the rat. Brain Res. 1994;655(1-2):241–245. doi:10.1016/0006-8993(94)91620-9

17. Walters MC, Sonner MJ, Myers JH, Ladle DR. Calcium imaging of parvalbumin neurons in the dorsal root ganglia. eNeuro. 2019;6(4). doi:10.1523/ENEURO.0349-18.2019

18. Yang Q, Ramamurthy A, Lall S, et al. Independent replication of motor cortex and cervical spinal cord electrical stimulation to promote forelimb motor function after spinal cord injury in rats. Exp Neurol. 2019;320:112962. doi:10.1016/j.expneurol.2019.112962

19. Carmel JB, Berrol LJ, Brus-Ramer M, Martin JH. Chronic electrical stimulation of the intact corticospinal system after unilateral injury restores skilled locomotor control and promotes spinal axon outgrowth. J Neurosci. 2010;30(32):10918–10926. doi:10.1523/JNEUROSCI.1435-10.2010

20. Irvine K-A, Ferguson AR, Mitchell KD, Beattie SB, Beattie MS, Bresnahan JC. A novel method for assessing proximal and distal forelimb function in the rat: the Irvine, Beatties and Bresnahan (IBB) forelimb scale. J Vis Exp. 2010;(46). doi:10.3791/2246

21. Irvine K-A, Ferguson AR, Mitchell KD, et al. The irvine, beatties, and bresnahan (IBB) forelimb recovery scale: an assessment of reliability and validity. Front Neurol. 2014;5:116. doi:10.3389/fneur.2014.00116

22. Carmel JB, Martin JH. Motor cortex electrical stimulation augments sprouting of the corticospinal tract and promotes recovery of motor function. Front Integr Neurosci. 2014;8:51. doi:10.3389/fnint.2014.00051

23. Metz GA, Whishaw IQ. The ladder rung walking task: a scoring system and its practical application. J Vis Exp. 2009;(28). doi:10.3791/1204

24. Sotocinal SG, Sorge RE, Zaloum A, et al. The Rat Grimace Scale: a partially automated method for quantifying pain in the laboratory rat via facial expressions. Mol Pain. 2011;7:55. doi:10.1186/1744-8069-7-55

25. Tan AM, Chakrabarty S, Kimura H, Martin JH. Selective corticospinal tract injury in the rat induces primary afferent fiber sprouting in the spinal cord and hyperreflexia. J Neurosci. 2012;32(37):12896–12908. doi:10.1523/JNEUROSCI.6451-11.2012

26. Kathe C, Hutson TH, McMahon SB, Moon LDF. Intramuscular Neurotrophin-3 normalizes low threshold spinal reflexes, reduces spasms and improves mobility after bilateral corticospinal tract injury in rats. Elife. 2016;5. doi:10.7554/eLife.18146

27. Mekhael W, Begum S, Samaddar S, et al. Repeated anodal trans-spinal direct current stimulation results in long-term reduction of spasticity in mice with spinal cord injury. J Physiol (Lond). 2019;597(8):2201–2223. doi:10.1113/JP276952

28. Carmel JB, Kimura H, Berrol LJ, Martin JH. Motor cortex electrical stimulation promotes axon outgrowth to brain stem and spinal targets that control the forelimb impaired by unilateral corticospinal injury. Eur J Neurosci. 2013;37(7):1090–1102. doi:10.1111/ejn.12119

29. Harvey P, Gong B, Rossomando AJ, Frank E. Topographically specific regeneration of sensory axons in the spinal cord. Proc Natl Acad Sci U S A. 2010;107(25):11585–90. doi: 10.1073/pnas.1003287107.

30. Asante CO, Martin JH. Differential joint-specific corticospinal tract projections within the cervical enlargement. PLoS One. 2013;8(9):e74454. doi: 10.1371/journal.pone.0074454.

